# TFAM-dependent mitochondrial integrity in regulatory T cells preserves immune landscape and restrains inflammaging

**DOI:** 10.1101/2022.03.08.483517

**Authors:** Kai Guo, Zhihan Wang, Jappreet Singh Gill, Trishna Debnath, Benu Bansal, Rayansh Poojary, Jacquiline Kim Correa, Hakan Celik, Het Mehta, Eden Abrham, Zachery Even, Mansib Rahman, Abby Lund Da Costa, Shilpi Jain, Xusheng Wang, Gopal Murugaiyan, Adrian T. Ting, Junguk Hur, Nadeem Khan, Holly Brown-Borg, Donald A Jurivich, Ramkumar Mathur

**Affiliations:** Department of Neurology, University of Michigan, Ann Arbor, MI 48109, USA; Department of Biomedical Sciences, School of Medicine and Health Sciences, University of North Dakota, Grand Forks, ND 58202, USA; Department of Geriatrics, School of Medicine and Health Sciences, University of North Dakota, Grand Forks, ND 58202, USA; West China School of Basic Medical Sciences & Forensic Medicine, Sichuan University, Chengdu, Sichuan, China; Department of Neurology, University of Tennessee Health Science Center, Memphis, TN 38163, USA; Ann Romney Center for Neurologic Diseases, Brigham and Women’s Hospital, Boston and Harvard Medical School, Boston, MA 02115, USA; Department of Immunology, Mayo Clinic, Rochester, MN 55905, USA; Dept of Oral Biology, College of Dentistry, University of Florida, Gainesville, FL 32603, USA; Department of Biomedical Data Science, School of Applied Computational Sciences, Meharry Medical College, Nashville, TN 37208, USA

## Abstract

*Foxp3⁺* regulatory T cells (Tregs) maintain immune homeostasis, yet the process that preserves their stability during aging remain unclear. Mechanistic progress has been hindered by models that ablate Tregs or delete *Foxp3*, which induce acute autoimmunity and prevent longitudinal study of physiological regulatory drift. Here, we establish a dose-dependent mitochondrial framework that preserves Treg lineage survival while permitting gradual metabolic attenuation. Using Treg-restricted TFAM modulation, a complementary haploinsufficient model, and whole-spleen single-cell profiling. We identify lineage-selective immune remodeling characterized by contraction of naïve CD8⁺ and follicular B-cell pools, alteration of CD4⁺ states, expansion of activated Tregs, and emergence of neuroimmune stress linked transcriptional modules that parallel physiological aging. Mechanistically, mitochondrial insufficiency is associated with functional loss of *FOXP3-*centered chromatin coordination and enrichment of NF-κB/NFAT/AP-1 inflammatory and senescence programs while lineage identity remains detectable. Partial mitochondrial attenuation within Tregs alone is sufficient to drive chronic low-grade systemic inflammation, neuromuscular decline, gut microbial restructuring, and elevated microglial responsiveness without Treg depletion. Pharmacologic and microbiota-directed interventions partially reduce inflammatory tone and improve functional metrics. Together, our findings identify TFAM as a key regulator of immune aging and reveal that healthy mitochondrial function in Tregs is essential for protecting against inflammaging and age-associated functional decline.

## Introduction

Immune aging is increasingly defined by dysregulated inflammatory tone instead of a uniform reduction in leukocyte counts^1, 2, 3, 4^. Converging evidence from human cohorts and murine models indicates that subtle cytokine elevation, redistribution of lymphoid subsets, and gradual functional drift occur prior to the manifestation of overt immunodeficiency, suggesting that the aging immune system is actively recalibrated rather than passively depleted. The gradual increase of long-lasting, low-grade inflammation is often called inflammaging which reflects disturbed immune balance rather than loss of immune cells, suggesting that regulatory pathways remain present but slowly change in function over time.

*Foxp3⁺* regulatory T cells are a metabolically and transcriptionally distinct lineage that rises predominantly within the CD4⁺ T cell compartment and enforce peripheral tolerance and suppress immunopathology^5, 6, 7, 8, 9^. In contrast, conventional effector T cells promote inflammation and primarily utilize glycolysis to support clonal expansion^10, 11^. The metabolic state of Tregs influences their epigenetic landscape, particularly at the conserved non-coding sequence 2 (CNS2) enhancer of the Foxp3 gene. Oxidative phosphorylation (OXPHOS) and fatty acid oxidation (FAO) support the maintenance of an open chromatin configuration at CNS2, facilitating sustained Foxp3 expression^12, 13, 14, 15^. This epigenetic regulation is essential for preserving Treg identity and suppressive function. Disruption of mitochondrial metabolism in Tregs can lead to decreased *Foxp3* expression and increased susceptibility to acquisition of effector-like programs. Specifically, impaired OXPHOS and FAO may result in the upregulation of transcription factors such as T-bet (TBX21), RORγt (RORC), and IRF4, which are associated with effector T cell lineages. This shift undermines Treg stability and promotes inflammatory responses. Emerging evidence, including from our own investigations, supports the concept that mitochondrial dysfunction is not merely correlative, but acts as a proximal driver of Treg instability, fueling systemic inflammation and age-associated immune remodeling^12, 16, 17, 18^. Progress in dissecting these mechanisms has been constrained by experimental systems that eliminate Tregs entirely, including scurfy mutations and diphtheria-toxin based *Foxp3* depletion models, which provoke rapid systemic autoimmunity and early lethality ^19, 20, 21^. Although these approaches firmly establish the necessity of *Foxp3* immune tolerance, they do not reproduce the gradual metabolic and transcriptional drift observed in physiological aging, where peripheral mature Tregs persist but slowly alter function. As a result, distinguishing intrinsic regulatory adaptation from secondary inflammatory collapse has remained challenging.

In recent years, mitochondrial genome integrity has emerged as an intrinsic checkpoint linking cellular metabolism to immune longevity^9, 22, 23^. Mitochondrial transcription factor A (TFAM), a nuclear-encoded high-mobility group (HMG) protein, is crucial for maintaining mitochondrial DNA (mtDNA) integrity and regulating redox homeostasis^24, 25^. Beyond its structural role in mtDNA packaging, TFAM modulates electron transport chain (ETC) function and limits mitochondrial reactive oxygen species (ROS) production. Loss of TFAM expression leads to mtDNA depletion, destabilization of oxidative phosphorylation (OXPHOS) complexes (CI-CIV), and activation of mitonuclear stress responses These include the mitochondrial unfolded protein response (UPR^mt^), integrated stress response (ISR) via ATF4/DDIT3, and downstream inflammatory signaling involving NF-κB and STAT1^25, 26, 27, 28^. The mitochondrial stress responses influence chromatin and transcriptional regulators, including histone deacetylases (HDACs), SIRT1, ten-eleven translocation (TET) enzymes, and polycomb repressive complex 2 (PRC2) components like EZH2. These regulators, which modulate histone acetylation/methylation and DNA demethylation states, are critical for *Foxp3* expression and the fidelity of the Treg lineage. Thus, redox-sensitive epigenetic rewiring under mitochondrial stress may mechanistically compromise Treg stability and function. However, whether TFAM-dependent mitochondrial integrity operates as an upstream checkpoint safeguarding immune homeostasis remains an unresolved question.

Interestingly, *Tfam* disruption in CD4⁺ T-cell compartment has been shown to accelerate systemic inflammation and tissue decline^17^. Prior studies employing Foxp3-directed TFAM deletion further demonstrate that mitochondrial impairment in regulatory T cells can precipitate acute inflammatory pathology or alter tumor progression^12, 16^. However, whether declining mitochondrial integrity confined specifically to *Foxp3⁺* Tregs is sufficient to reorganize whole-immune architecture toward an aging-like trajectory rather than merely reproducing inflammatory syndromes associated with overt regulatory dysfunction has remained unclear. Our single-cell transcriptomic analyses revealed that *FOXP3⁺* T cells exhibit mitochondrial stress–associated signatures that track with inflammatory skewing in human aging and frailty, implicating mitochondrial dysfunction in progressive immune remodeling^18^. Consistent patterns reported in independent cohorts further strengthen the association between mitochondrial perturbation and age-related shifts in Treg state and function^30, 31, 32, 29^. Collectively, these single-cell studies suggest that early mitochondrial dysfunction could involve in the gradual re-shaping of immune trajectories.

Here, to dissect this relationship, we employed two Treg-specific TFAM mouse models: a conditional knockout (Foxp3⁺ᶜʳᵉTfamᶠ^l^/ᶠ^l^) and a heterozygous model (Foxp3⁺ᶜʳᵉTfam^ᶠ/+)^, which exhibits a moderate physiological reduction in TFAM expression. Together, these models recapitulate aspects of the progressive mitochondrial decline observed during both normal and pathological aging. Mitochondrial dysfunction in the Treg lineage led to a cascade of immunological issues, including hyper-inflammatory CD4^+^ T cell fate changes and altered gene expression. Such shifts are consistent with hallmarks of inflammaging, which include chronic low-grade inflammation, gut microbial dysbiosis, and neuroimmune activation. Together, our study establishes a scalable experimental framework to directly examine how gradual mitochondrial decline in persistent Tregs alters systemic inflammatory set-points. Our findings establish mitochondrial integrity not only as a metabolic feature but also as a modifiable regulator of Treg quality and immune homeostasis, highlighting mitochondrial maintenance as a potential avenue to mitigate chronic age-associated inflammation.

## Results

### Treg-intrinsic TFAM loss in Foxp3⁺ Tregs reprograms CD4⁺ T cell lineage trajectories and enforces effector bias

To investigate how mitochondrial TFAM deficiency in *Foxp3⁺* Tregs perturb systemic CD4⁺ T cell homeostasis, we conducted single-cell RNA sequencing (scRNA-seq) on whole spleens from Foxp3^Cre^Tfam^f/f^ (TFAM cKO) and Tfam^f/f^ (wild-type) (Fig. 1A). This model selectively impairs mitochondrial DNA maintenance in mature Tregs while largely preserving Foxp3 expression and lineage presence, enabling assessment of regulatory *functional* dysfunction rather than the severe autoimmune phenotypes produced by Foxp3-null/scurfy or diphtheria-toxin depletion systems ^19, 20, 21^. Following stringent quality control, we retained transcriptomic profiles from 14,198 cells in TFAM cKO and 34,958 cells in wild-type mice, excluding erythroid lineage cells from downstream analyses (Fig. 1B). Based on canonical immune markers, we resolved 21 major immune subsets across 26 transcriptionally distinct clusters conserved across both genotypes (Fig. 1B; Fig. S1A-C). Notably, naïve-associated CD8⁺ T cells marked by Tcf7 and Lef1 were reduced, as were follicular B-cell populations expressing Cd79b and Fcer2a, whereas total CD4⁺ abundance remained relatively stable but underwent pronounced transcriptional redistribution. Importantly, the contraction of naïve CD8⁺ cells was not interpreted as direct mitochondrial impairment within CD8⁺ cells, but rather as a secondary consequence of altered regulatory set-points consistent with physiological immune aging in which naïve CD8⁺ attrition is a prominent and early feature of compartmental drift ^1, 2^.

**Figure 1.**
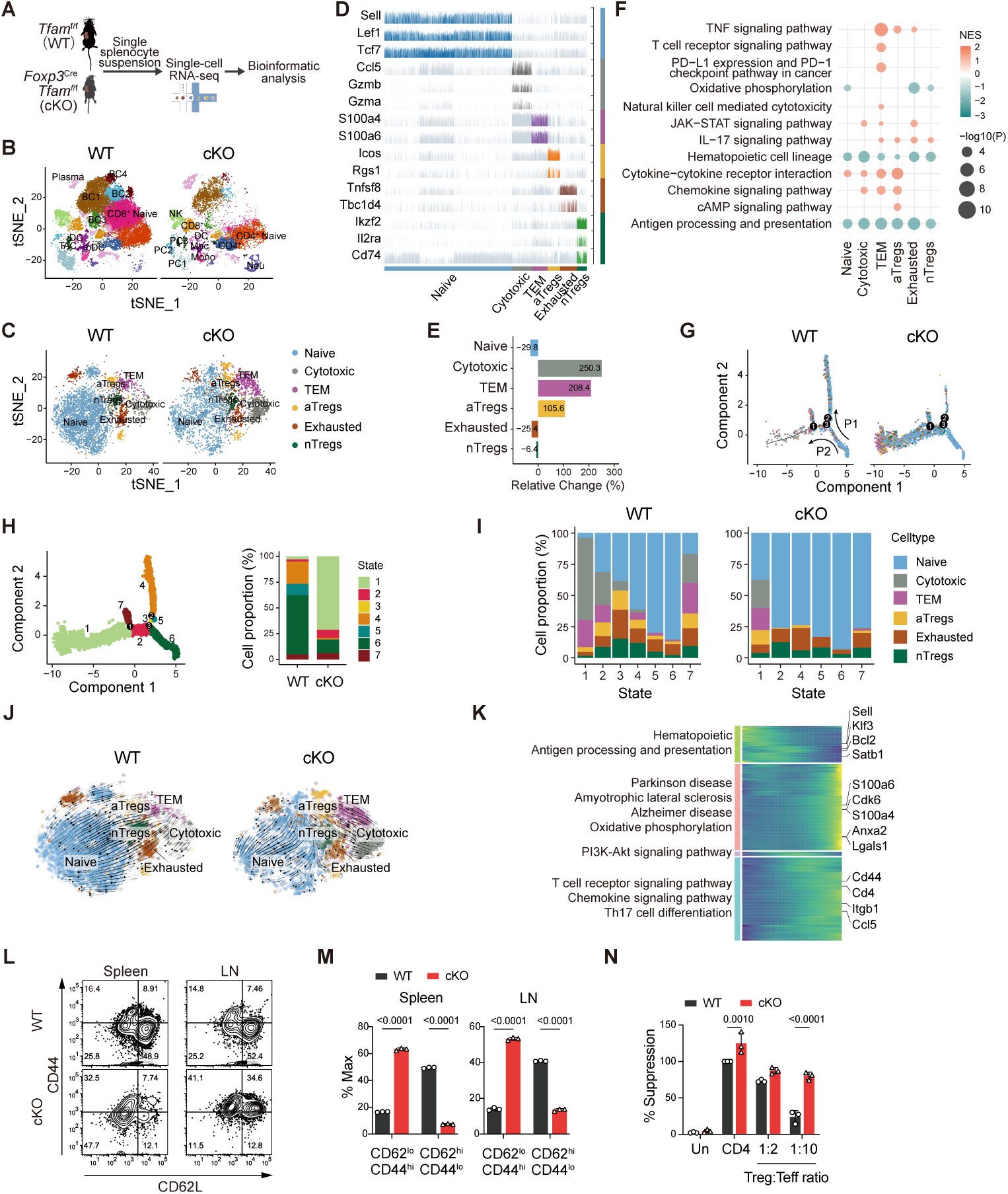
Treg-intrinsic TFAM loss reprograms CD4⁺ T cell lineage fate toward pro-inflammatory and terminal effector states. **(A)** Schematic representation of the experimental workflow. Splenocytes were isolated from Foxp3^Cre^Tfam^f/f^ (TFAM cKO) and Tfam^f/f^ (WT) mice for single-cell RNA sequencing (scRNA-seq) and downstream analysis. **(B)** UMAP projection of 49,156 immune cells (WT: 34,958; cKO: 14,198), identifying 21 major immune subsets across 26 transcriptional clusters. Erythroid lineage cells were excluded. **(C)** UMAP projection of CD4⁺ T cells (WT: 4,324; cKO: 3,497), revealing six transcriptionally distinct subsets: naïve, effector memory (TEM), exhausted, natural Tregs (nTregs), aged/dysfunctional Tregs (aTregs), and cytotoxic CD4⁺ T cells. **(D)** Track plot showing representative gene expression patterns across single CD4⁺ T cells. Rows represent genes; columns represent individual cells. (**E)** Bar graph quantifying relative abundance of CD4⁺ T cell subsets in WT and cKO mice. TFAM deficiency promotes expansion of effector-like states (aTregs, cytotoxic, and TEM), with reduced naïve and exhausted populations. (**F**) Gene set enrichment analysis (GSEA) of KEGG pathways shows increased TNFα signaling, cytokine/chemokine responses, and calcium flux in CD4⁺ T cells from TFAM cKO mice. (**G**) Pseudotime trajectory analysis (Monocle) of CD4⁺ T cells reveal two dominant differentiation paths (P1 and P2). TFAM-deficient cells preferentially adopt the alternative P2 trajectory, enriched in transcriptional States 1 and 2. (**H**) Monocle 2 trajectory cell states for all CD4^+^ T cells, each branch represents one cell state (left). Barplots representing cell proportions in the seven states of WT and cKO (right). Cells are colored according to seven states, which partition the trajectory. (**I**). Relative proportion of CD4^+^ T cells among different states of WT and cKO spleen, color stands for cell type. (**J**) RNA velocity analysis confirms directionality of lineage bifurcation, with streamlines overlaid on UMAP embedding indicating transcriptional flow. (**K**) Heatmap of four gene clusters ordered by pseudotime. Cluster 1 genes (e.g., Klf3, Bcl2) mark early developmental states; Cluster 2 genes (e.g., Ccl5, S100a6) are linked to terminal effector function and inflammation. KEGG pathway enrichment of cluster-specific genes is shown. (**L**) Representative flow cytometry plots of CD4⁺CD44^hi^CD62L^lo^ effector and CD4⁺CD44^lo^CD62L^hi^ naïve T cells in spleen and lymph nodes from WT and TFAM cKO mice. (**M**) Quantification of mean fluorescence intensity (MFI) for effector and naïve CD4⁺ T cells across organs. TFAM cKO mice exhibit increased effector T cell signatures. (**N**) Suppression assay showing reduced regulatory function of TFAM-deficient Tregs in vitro. Co-culture with CFSE-labeled CD4⁺ Tconv cells demonstrates impaired suppression of proliferation upon CD3/CD28 stimulation.

TFAM deletion in Tregs elicited a striking inflammatory reprogramming of the CD4⁺ T cell compartment, marked by enrichment of TNF signaling, inflammation-associated aging pathways, and gene signatures linked to tumorigenic transformation (Fig. S1D; Table S1). Upregulation of pro-inflammatory mediators including *S100a6* and *Zfp3612*, coupled with diminished expression of homeostatic regulators such as *Cd79b* and *Fcer2a*, was consistent with premature immune aging (Fig. S1E). To examine how TFAM-deficient Tregs influence CD4⁺ T cell diversity at a steady state, we profiled subset heterogeneity based on canonical markers. Based on canonical lineage-defining markers, six transcriptionally defined CD4⁺ T cell subsets were identified: naïve (*Sell, Lef1, Tcf7*), effector memory (TEM; *S100a4, S100a6*), exhausted (*Tnfsf8, Tbc1d4*), conventional Tregs (nTregs; *Ikzf2, Il2ra, Cd74*), activated Tregs (aTregs; *Icos, Rgs1*), and cytotoxic CD4⁺ T cells (*Ccl5, Gzma, Gzmb*) (Fig. 1C, D). Comparative analysis revealed substantial shifts in subset proportions in TFAM cKO mice (Fig. 1E, Fig. S2A), including marked expansion of aTregs (+105.6%), cytotoxic CD4⁺ T cells (+250.3%), and TEM cells (+208.4%), alongside reduced frequencies of exhausted (-25.4%) and naïve (-29.8%) subsets, and a modest reduction in nTregs (-6.4%). These shifts reflect an altered balance favoring pro-inflammatory and terminally differentiated effector states. Thus, TFAM attenuation preserved lineage identity but reshaped qualitative fate composition, aligning with aging datasets in which Tregs persist yet adopt activated, cytokine-responsive phenotypes rather than disappearing ^30, 31^.

To gain mechanistic insights into these phenotypic alterations, differential gene expression and pathway enrichment analyses in CD4⁺ T cells from TFAM cKO versus WT mice uncovered strong upregulation of TNFα signaling, calcium flux, and cytokine/chemokine networks, with concurrent downregulation of hematopoietic lineage maintenance programs (Fig. 1F, Fig. S2B, Table S2). IL-17 signaling was particularly enriched across TEM, exhausted, aTregs, and nTregs, suggesting a bias toward pro-inflammatory Th17-like conversion a hallmark of increased Treg plasticity. Recognizing that CD4⁺ T cell diversity is governed by dynamic lineage trajectories, we next applied trajectory inference to map differentiation states and fate decisions within CD4⁺ T cell compartments. Trajectory inference using Monocle revealed a branched differentiation architecture with three bifurcation points and two dominant paths (P1 and P2), covering seven transcriptional states (Fig. 1G-I, Fig. S2C-D). CD4⁺ T cells from WT mice primarily followed the P1 trajectory, occupying States 4-6, whereas TFAM cKO cells were enriched in States 1, 2, and 6, indicating a skewing toward the P2 trajectory and altered fate commitment. Naïve T cells, while concentrated in States 4-6 in both groups, were aberrantly distributed across States 1, 2, and 7 in cKO mice. Notably, aTregs were enriched in State 1, while cytotoxic CD4⁺ T cells were depleted in terminal States 1 and 7 (Fig. 1I). The shifts point to preferential lineage commitment along the alternative P2 trajectory in TFAM-deficient cells, contrasting with the homeostatic P1 path favored by wild-type counterparts. Next, we applied RNA velocity analysis to confirm the inferred directionality of fate transitions, supporting Monocle-inferred pseudotime dynamics (Fig. 1J). Gene expression kinetics along pseudotime identified four temporally distinct clusters. Early expressed Cluster 1 genes (*Klf3, Sell, Bcl2, Satb1*) were enriched in hematopoietic progenitors, while Cluster 2 genes (*Anxa2, S100a4, Ccl5, S100a6, Lgals1*) were associated with terminal effector differentiation and inflammation (Fig. 1K). Strikingly, Cluster 2 was enriched for gene signatures linked to neurodegenerative diseases, Alzheimer’s, Parkinson’s, and ALS, as well as oxidative phosphorylation (Fig. 1K, Table S3). KEGG analysis across Monocle states further demonstrated that State 1, enriched in TFAM-deficient cells, was associated with neuroinflammatory and neurodegeneration-related programs, whereas hematopoietic renewal pathways dominated the WT-favored P1 trajectory (Fig. 1K, Fig. S2D, Table S4). These findings suggest that Treg mitochondrial dysfunction promotes effector CD4⁺ T cell programs enriched for transcriptional features observed in neuroinflammatory and neurodegenerative conditions. Flow cytometry confirmed the expansion of CD4⁺CD44^hi^CD62L^lo^ effector T cells and depletion of CD4⁺CD44^lo^CD62L^hi^ naïve populations in secondary lymphoid organs of TFAM cKO mice (Fig. 1L-M). Functionally, TFAM-deficient Tregs exhibited reduced suppressive capacity, allowing for enhanced proliferation of wild-type responder CD4⁺ T cells in co-culture assays (Fig. 1N). Collectively, these findings indicate that mitochondrial transcriptional attenuation restricted to Foxp3⁺ Tregs associates with coordinated splenic immune remodeling characterized by naïve CD8⁺ contraction, follicular B-cell reduction, diversification of CD4⁺ states, selective expansion of ICOS⁺ activated Tregs, and enrichment of stress- and IL-17–linked transcriptional programs. In aggregate, the resulting immune landscape aligns with aging-like immune drift redistribution and functional reprogramming across compartments rather than a depletion-equivalent phenotype or synchronized effector expansion typical of acute Treg ablation.

### TFAM loss in Tregs unleashes a multi-lineage stress response coupled with inflammatory and senescence signatures

To investigate how mitochondrial TFAM deficiency disrupts transcriptional networks in Foxp3⁺ Tregs, particularly those governing inflammation-associated regulons and downstream TNF-driven signaling implicated in Treg destabilization and effector skewing of CD4⁺ T cells, we performed gene regulatory network (GRN) inference using scRNA-seq data from TFAM cKO and wild-type spleens. We identified 177 high-confidence regulons distributed across CD4⁺ T cell subsets (Fig. 2A). Canonical immunoregulatory transcription factors, including *Irf2, Irf8, Irf9,* and *Bcl11a*, were preferentially enriched in wild-type Tregs, reflecting preservation of suppressive lineage identity. In contrast, TFAM-deficient Tregs showed marked activation of inflammatory regulons such as *Nfkb2, Nfatc1, Rora,* and *Runx2*, consistent with a breach in the mitochondrial checkpoint that permits effector conversion (Fig. 2A, Fig. S3). Stress-adaptive regulons, including *Egr1, Junb,* and *Xbp1*, were enriched in exhausted T cells, aTregs, and even nTregs in TFAM cKO mice, pointing to a convergent transcriptional stress response. Enrichment of *Runx3* in cytotoxic CD4⁺ subsets further highlighted mitochondrial TFAM as a key repressor of cytolytic plasticity. Notably, upregulation of *Ddit3* across all CD4⁺ subsets indicated a shared mitochondrial stress program (Fig. 2A). Targeted expression profiling of inflammatory regulators, including *Prdm1, Nfkb1, Nfkb2,* and *Ahr*, showed strong induction in TFAM-deficient Tregs, reinforcing the destabilization of the regulatory axis (Fig. 2B). The ensuing Systemic inflammation further activates myeloid-derived cells, such as monocytes and macrophages, enhancing their oxidative phosphorylation activity. This heightened metabolic state contributes to increased neuroinflammation, and the activation of neurodegenerative pathways associated with Parkinson’s and Alzheimer’s diseases (Fig. 2C). To assess downstream consequences, bulk RNA-seq of TFAM cKO splenocytes revealed strong induction of cell death pathways, including apoptosis and necroptosis (Fig. 2D). Immunoblotting confirmed caspase-3 and -8 cleavage, IκBα degradation, and phosphorylation of RIPK1, RIPK3, and MLKL signatures of RIPK-mediated immunogenic cell death (Fig. 2E). This was substantiated in situ by Annexin V/PI staining, which revealed marked increases in apoptotic and necroptotic cell populations in both splenic and cutaneous tissues (Fig. S4E), indicating systemic cytopathology secondary to mitochondrial collapse in Tregs. Collectively, these findings lend support to regulatory destabilization caused by chronic inflammatory injury rather than wholesale Treg lineage elimination.

**Figure 2.**
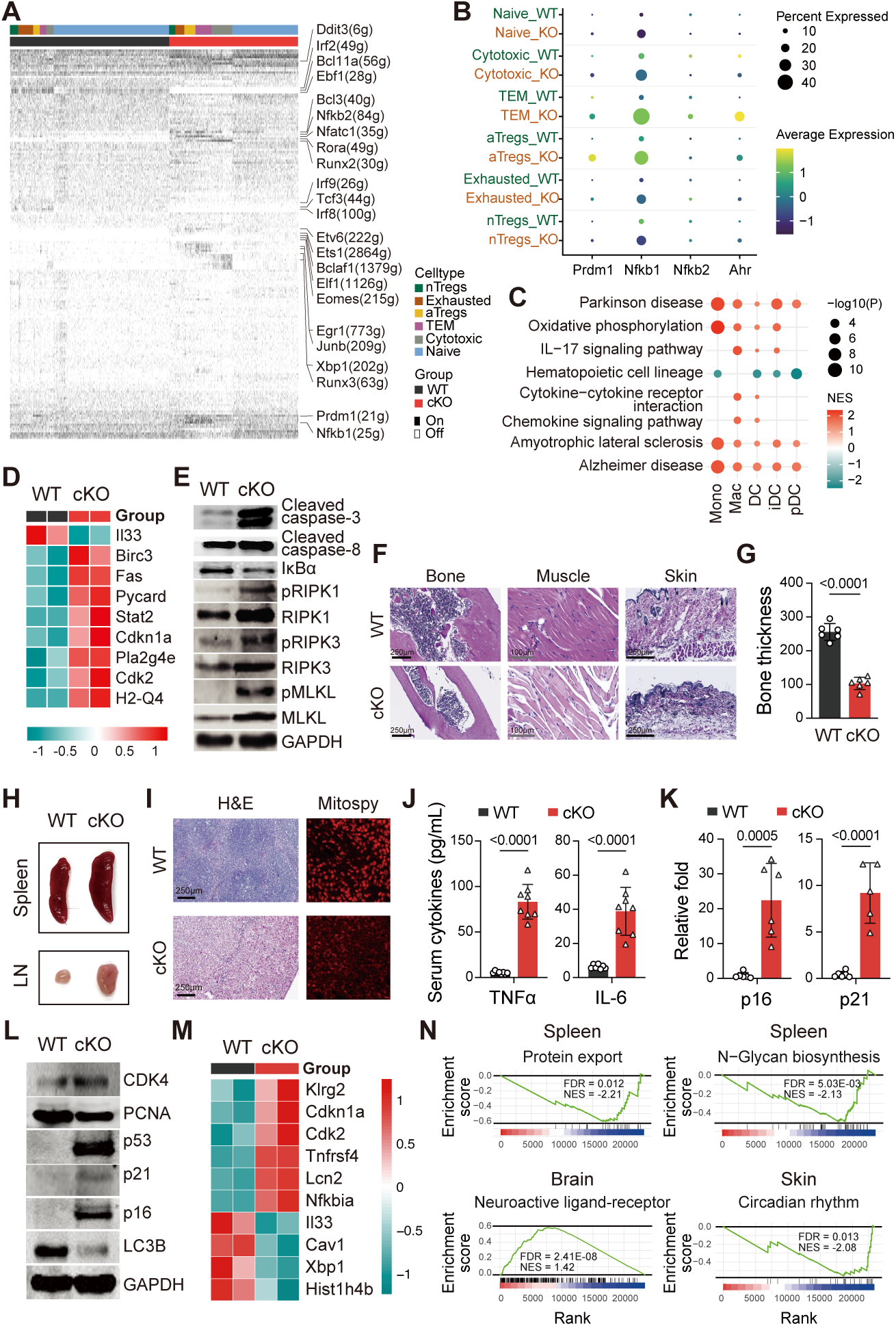
Treg-intrinsic TFAM loss triggers a multi-lineage stress response, inflammatory reprogramming, and systemic senescence. (**A**) Gene regulatory network (GRN) inference from scRNA-seq data identified 177 regulons across CD4⁺ T cell subsets. TFAM cKO Tregs showed increased inflammatory activity (Nfkb2, Nfatc1, Rora, Runx2), stress-responsive (Egr1, Junb, Xbp1), and cytotoxic (Runx3) regulons, alongside pan-CD4⁺ expression of the mitochondrial stress gene Ddit3. (**B**) Dot plots showing upregulation of key inflammatory transcriptional regulators (Prdm1, Nfkb1, Nfkb2, Ahr) in TFAM-deficient Tregs. (**C**) Gene set enrichment analysis (GSEA) shows significant pathways in Monocytes (Mono), Macrophages (Mac), dendritic cell (DC), Immature Dendritic Cells (iDC) and Plasmacytoid Dendritic Cells (pDC) cells in TFAM cKO mice. (**D**) Bulk RNA-seq of cKO spleens revealed enrichment of cell death pathways, including apoptosis and necroptosis. (**E**) Immunoblots confirmed caspase-3 and -8 cleavage, IκBα degradation, and phosphorylation of RIPK1, RIPK3, and MLKL, indicative of RIPK-mediated immunogenic cell death. (**F**) Histological analysis revealed multi-organ inflammatory infiltration in colon, liver, lung, and kidney of TFAM cKO mice. (**G**) H&E staining of femurs showed pronounced cortical bone thinning in TFAM cKO animals. (**H**) Spleens from TFAM cKO mice exhibited splenomegaly, follicular disruption, and marginal zone disorganization. (**I**) Mitospy staining in sorted Tregs revealed mitochondrial depolarization and elevated oxidative stress in TFAM-deficient cells. (**J**) Elevated circulating levels of TNF-α, IL-6, and IFNγ in serum from TFAM cKO mice, confirming systemic inflammation. (**K**) RT-qPCR showed increased expression of Cdkn2a (p16) and Cdkn1a (p21); (**L**) Immunoblotting of splenic lysates confirmed elevated levels of p53, CDK4, PCNA, p16, and p21, consistent with senescence-associated gene expression. (**M**) Whole-spleen RNA-seq revealed enrichment of aging- and inflammation-associated gene networks. (**N**) GSEA showed significant upregulation of pathways linked to neuroactive ligand–receptor signaling, and downregulation of circadian rhythm disruption, and protein trafficking which are hallmarks of systemic immune aging.

To determine if Treg-specific mitochondrial dysfunction initiates systemic decline, we monitored TFAM cKO mice longitudinally. While born at Mendelian ratios, these mice developed progressive organismal deterioration from 4 weeks, including alopecia, kyphosis, lymphoid organomegaly, impaired mobility, and early mortality (p<0.0001; Fig. S4A-B). By 8-10 weeks, they developed severe skeletal muscle atrophy, adipose tissue loss, cortical bone thinning, and markedly reduced survival relative to wild-type littermates (Fig. S4C). Histological analysis revealed inflammatory infiltration in the lung, liver, colon, and kidney (Fig. 2F, S5A), and cortical bone loss (Fig. 2F, 2G). Spleens from TFAM cKO mice exhibited splenomegaly, marginal zone disruption, and follicular disorganization (Fig. 2H). These phenotypes were consistent with disrupted immune regulation downstream of Treg-intrinsic mitochondrial dysfunction. Mitospy analysis showed mitochondrial depolarization and increased oxidative stress in cKO Tregs (Fig. 2I, Fig. S4D, S4E). Multiplex cytokine profiling revealed a pronounced pro-inflammatory signature characterized by elevated levels of IL-17, IFN-γ, IL-12, IFN-β, GM-CSF, and IL-27, in parallel with significantly increased circulating TNF-α and IL-6, reflecting systemic immune activation and inflammatory amplification (Fig. 2J; Fig. S5B). At the molecular level, cKO mice exhibited upregulation of *Cdkn2a (p16)* and *Cdkn1a (p21)* (Fig. 2K), and immunoblots confirmed elevated p53, CDK4, p16, and p21 (Fig. 2L), indicative of a senescence-like program. Whole-spleen RNA-seq revealed enrichment of inflammatory and aging-related networks (Fig. 2M). Gene Set Enrichment Analysis (GSEA) from spleen, skin and brain revealed enrichment of neuroactive signaling, circadian disruption, and protein trafficking defects pathways associated with immune aging and systemic decline (Fig. 2N, Table S5). These findings establish TFAM as a critical mitochondrial checkpoint that safeguards Treg identity, restrains inflammation, and maintains immune equilibrium.

### Partial TFAM reduction in Tregs as a critical driver of aging pathologies, neuroinflammation, and cognitive decline

To model the progressive mitochondrial decline characteristic of physiological aging, we transitioned to a Treg-specific TFAM heterozygous mouse (TFAM Het; *Foxp3^Cre^TFAM^f/+^*). While complete TFAM ablation in Tregs provides mechanistic insight into the necessity of mitochondrial integrity for immune homeostasis, the rapid-onset pathology of this model precludes longitudinal interrogation of chronic immune aging and neurodegenerative processes. To better understand, the TFAM Het model, where one functional allele of the *Tfam* gene is deleted specifically in CD4⁺Foxp3⁺ Tregs, resulting in a sustained ∼50% reduction in TFAM expression in CD4⁺Foxp3⁺ Tregs. This genetically defined model recapitulates subclinical mitochondrial insufficiency and mirrors the progressive mitochondrial decline characteristic of the human aging trajectory. Supporting its translational relevance, analysis of human PBMC transcriptomes revealed a significant age-dependent decline in TFAM expression, reinforcing its role as a mitochondrial-immune rheostat (Fig. 3A). By 8-10 months, TFAM Het mice exhibited classic hallmarks of aging, including kyphosis, weight loss, and reproductive decline (Fig. 3B-C, Fig. S6A-B).

**Figure 3.**
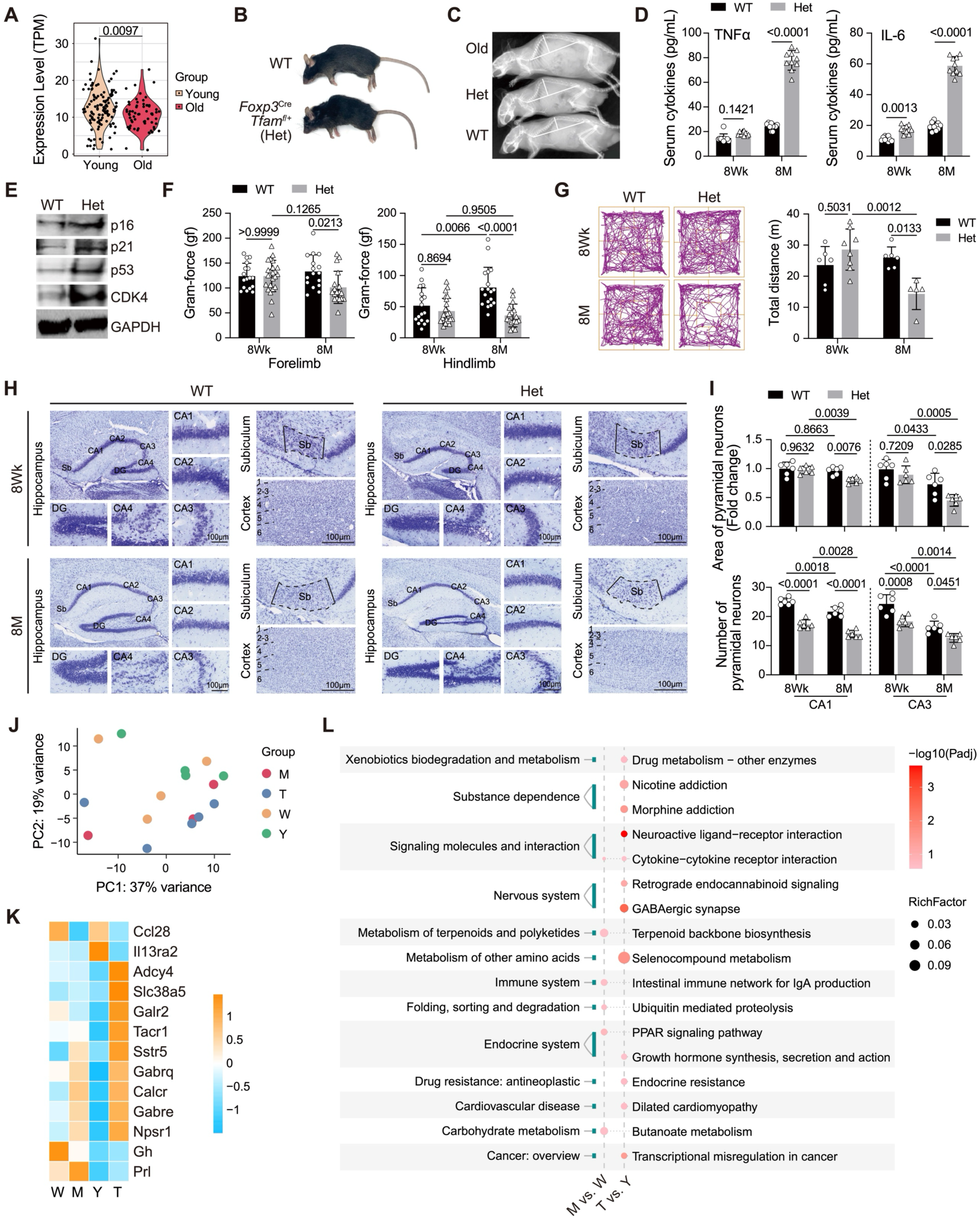
Partial TFAM reduction in Tregs promotes systemic aging features, peripheral inflammation, and early neurodegenerative changes. (**A**) Human PBMC transcriptome analysis reveals age-associated decline in TFAM expression, supporting its role as a mitochondrial-immune rheostat. (**B-C**) TFAM Het (Foxp3^Cre^Tfam^f/+^) mice develop aging phenotypes, including kyphosis and reduced body weight, by 8-10 months of age. (**D**) Circulating TNF-α and IL-6 levels are elevated in TFAM Het mice, indicating persistent low-grade systemic inflammation. (**E**) Splenic CD4⁺ T cells isolated from TFAM Het mice display elevated expression of senescence-associated markers, including Cdkn2a (*p16^Ink4a*), Cdkn1a (*p21^Cip1*), p53, and CDK4, with GAPDH serving as the loading control. (**F**) Neuromuscular assessment shows reduced forelimb grip strength in TFAM Het mice. (**G**) Open field behavioral assay demonstrates decreased exploratory behavior and locomotor hypoactivity, consistent with CNS dysfunction. (**H–I**) Cresyl violet staining reveals neuronal loss in the hippocampus and cortex of 8-month-old TFAM Het mice compared to 8-week controls. (**J**) Longitudinal hippocampal RNA-seq profiling shows minimal transcriptional changes at 8 weeks but extensive neuroimmune remodeling by 8 months. (**K**) Heatmap displaying upregulation of neuroinflammation-associated genes (Gabra5, Slc38a5, Calcr, Ganbare) in 8-month TFAM Het hippocampus. (**L**) KEGG pathway enrichment of 8-month hippocampal transcriptomes identify significant enriched pathways of oxidative stress, inflammatory signaling, and neurodegeneration-linked pathways.

Histopathological evaluation demonstrated that 8-month-old TFAM cKO mice exhibited pronounced structural degeneration in the spleen and lung, markedly exceeding that observed in both 8-week-old controls and TFAM Het counterparts. Notably, the extent of tissue deterioration in TFAM-deficient mice phenocopied the age-associated pathology typically seen in 2-year-old wild-type mice, underscoring the accelerated aging phenotype driven by Treg-intrinsic mitochondrial dysfunction (Fig. S6C). Senescence markers *Cdkn2a (p16^Ink4a^)* and *Cdkn1a (p21^Cip1^)* were significantly upregulated in splenic CD4⁺ T cells (Fig. 3D), implicating mitochondrial stress induced senescence. Circulating TNF-α and IL-6 levels were markedly elevated (Fig. 3E), indicative of persistent low-grade inflammation. Neuromuscular evaluation revealed diminished grip strength (Fig. 3F), and behavioral assays demonstrated reduced exploratory activity and locomotor hypoactivity (Fig. 3G), consistent with CNS dysfunction. To interrogate the neuroanatomical consequences of peripheral Treg mitochondrial instability, cresyl violet staining of brain sections revealed substantial neuronal loss in the hippocampus and cortex of 8-month-old TFAM Het mice relative to 8-week counterparts (Fig. 3H-I). PCA plots from longitudinal hippocampal RNA-seq profiling revealed a clear difference between Het8M group (T) and Het 8W group (Y) (Fig. 3J), Notably, compared to both 8-week-old wild-type and age-matched 8-month-old controls we observed marked upregulation of neural stress–associated genes including *Gabra5*, *Slc38a5*, *Calcr*, and *Ganbare* (Fig. 3K). Further compared to wild 8W and comparable 8M TFAM Het mice shown KEGG pathway enrichment shown a marked elevated, inflammatory signaling, Garabergic synapses neuro-active-ligand-receptor interaction and neurodegeneration-linked pathways (Fig. 3L). To validate the functional relevance of these molecular changes, we next assessed neuroimmune phenotypes at the cellular level. Iba1 immunohistochemistry revealed widespread activation of amoeboid microglia in TFAM Het brains (Fig. 4A-B), with NeuN co-staining confirming adjacent neuronal loss. Flow cytometric analysis revealed elevated MHC-I expression on brain-resident CX3CR1⁺ microglia in late-phase TFAM Het mice (Fig. 4C), indicating enhanced antigen presentation. Upon TNF-α stimulation, purified CX3CR1⁺ microglia from TFAM Het mice exhibited increased expression of IL-1β, TNFR1, and IL-6 transcripts, highlighting a heightened sensitivity to TNF-driven inflammatory activation compared to WT controls (Fig. 4C-D). These findings collectively support a peripheral-to-central immunometabolic cascade, wherein Treg-intrinsic mitochondrial insufficiency initiates systemic inflammation, senescence, and CNS immune activation. The TFAM Het model reveals that gradual mitochondrial decline in CD4⁺ Tregs is sufficient to disrupt systemic immune homeostasis, propagate neuroinflammation, and accelerate degenerative aging processes.

**Figure 4.**
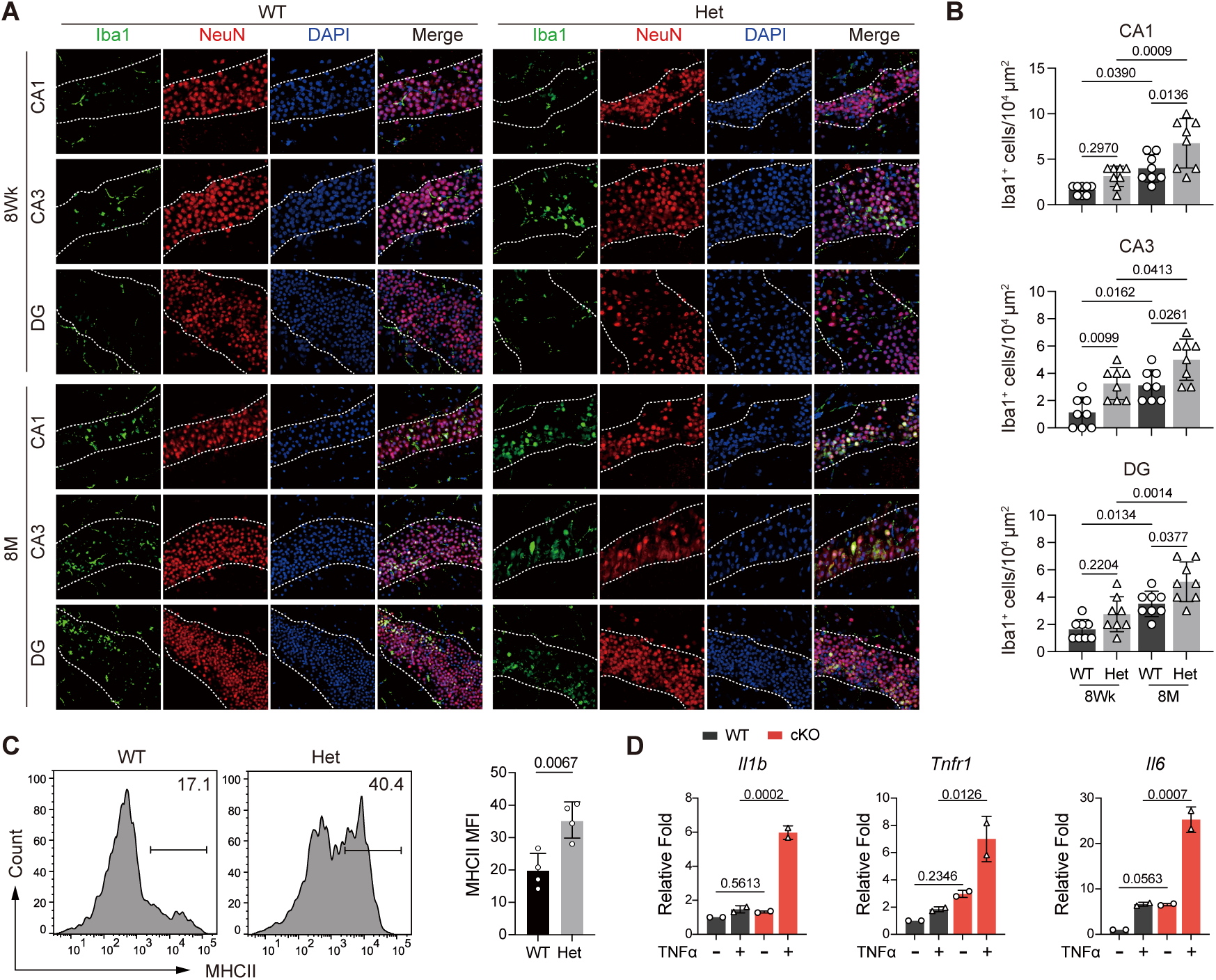
TFAM insufficiency in Tregs triggers CNS immune activation and microglial sensitization to TNF signaling. (**A-B**) Iba1 and NeuN immunohistochemistry shows widespread activation of amoeboid microglia in the cortex and hippocampus of 8-month-old TFAM Het mice, with concurrent neuronal loss. (**C**) Flow cytometric analysis reveals increased MHC-I expression on brain-resident CX3CR1⁺ microglia from TFAM Het mice, consistent with enhanced antigen presentation. (**D**) Ex vivo stimulation with TNF-α induces elevated *Il1b*, *Tnfr1*, and *Il6* expression in CX3CR1⁺ microglia from TFAM Het brains, indicating heightened inflammatory sensitivity.

### TFAM insufficiency in Tregs drives microbiota-lipid dysregulation and neuroinflammatory reprogramming via the gut-brain-immune axis

Mitochondrial metabolism in immune cells is closely coupled to host lipid flux and microbiota-derived metabolites^32^. We showed that advancing age is accompanied by progressive dysregulation of host–microbial interactions, forming a bidirectional cycle linking metabolism, microbial composition, and immune regulation^33^. Recently CD4 T cell therapy shown counteracts inflammaging and senescence by preserving gut barrier integrity^34^. We therefore asked whether gradual TFAM reduction within regulatory T cells also alters the intestinal metabolic environment and downstream immune tone. Stable Treg identity depends on intact oxidative metabolism. Short-chain fatty acids (SCFAs), especially butyrate and propionate, reinforce FOXP3 expression through mitochondrial–epigenetic coupling^35, 36, 37^. It has remained unclear whether long-lived Tregs undergoing slow mitochondrial decline can influence microbial ecology. Using a graded TFAM-insufficiency strategy that preserves lineage survival while attenuating mitochondrial transcription, and applying longitudinal 16S sequencing using our previously published protocol in aging mouse model ^33^, we observed progressive, age-associated restructuring of gut microbial communities in TFAM Het mice (Fig. 5A–B). Beta-diversity analyses separated groups by both genotype and age (Fig. 5B). This pattern indicates gradual community redirection rather than abrupt dysbiosis. At the phylum level, aged TFAM Het animals showed relative expansion of Campylobacterota and contraction of Bacteroidota (Fig. 5C). Shared-taxa analysis revealed a preserved core microbiota across age-matched groups (Fig. 5D). This overlap suggests that these specific bacterial changes are associated with aging processes common to both genotypes, highlighting potential age-related microbial shifts that occur irrespective of TFAM status.

**Figure 5.**
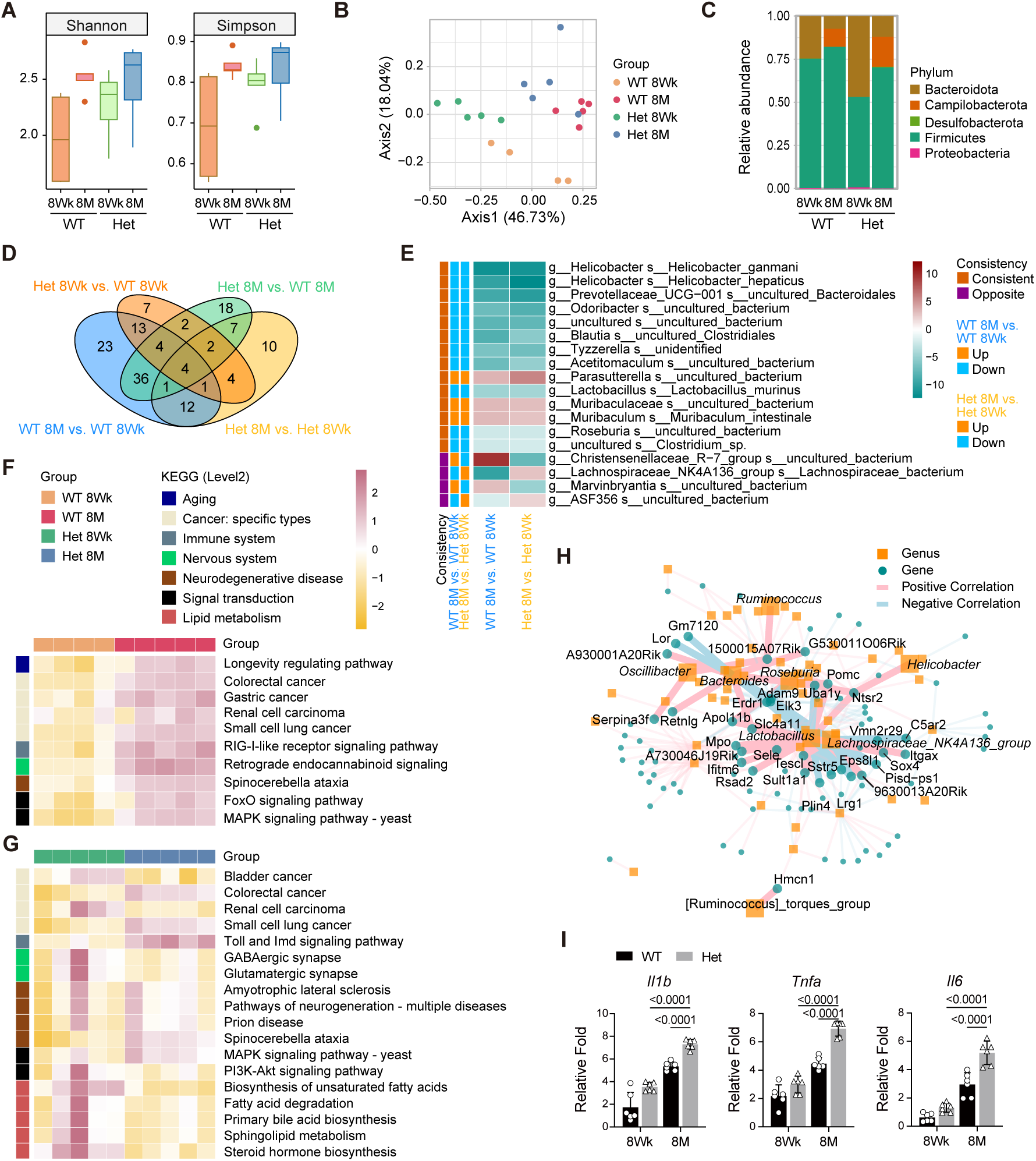
TFAM insufficiency in Tregs drives microbiota–lipid dysregulation and neuroinflammatory reprogramming via the gut–brain–immune axis. (**A**) Longitudinal 16S rRNA sequencing reveals significantly elevated α-diversity (Shannon and Simpson indices) in TFAM Het mice, peaking at 8 months of age. (**B**) β-diversity analysis (Bray–Curtis dissimilarity) demonstrates genotype- and age-specific microbial clustering, indicating progressive community remodeling. (**C**) Phylum-level taxonomic profiling reveals expansion of *Campilobacterota* and depletion of *Bacteroidota* in aged TFAM Het mice, signatures linked to inflammaging. (**D**) VennDiagram depicting distribution and overlap of microbial species across genotypes and age groups. (**E**) Differential abundance analysis identifies 18 significantly altered taxa, including loss of key SCFA-producing genera (*Lactobacillus murinus*, *Odoribacter*, *Prevotellaceae_UCG-001*) in TFAM Het mice. (**F**) KEGG-based predictive metagenomics shows TFAM Het microbiota enriched in age-related pathways including FoxO, and spinocerebellar ataxia signaling. (**G**) Lipid metabolism pathways, including unsaturated fatty acid biosynthesis, bile acid biosynthesis, and sphingolipid metabolism, are significantly altered. Neurotransmission pathways (GABAergic, glutamatergic synapses) are also enriched in aged TFAM Het mice. (**H**) Microbiota–host gene co-correlation network analysis reveals 323 significant interactions. Notably, *Helicobacter* positively correlates with *Ntsr2*, while Lactobacillus inversely correlates with neuroendocrine (e.g., *Pomc*, *Sstr5*) and developmental genes (*Elk3*, *Sox4*) and positively correlates with inflammatory genes (*Mpo*, *Ifitm6*). *Lachnospiraceae_NK4A136_group* links to complement-mediated inflammation (*C5ar2*). (**I**) RT-qPCR validation confirms elevated expression of *Il1b* and *Tnfr1* in peripheral tissues from TFAM Het mice, supporting a chronic inflammatory state.

Notably, differential abundance analysis identified 18 aging-associated microbial taxa altered across genotypes, including the depletion of canonical SCFA-producing lineages, such as *Lactobacillus murinus*, *Odoribacter*, and *Prevotellaceae_UCG-001* (Fig. 5E). Given that SCFAs like butyrate are critical for Treg stability, intestinal barrier integrity, and peripheral immune tolerance, these compositional shifts suggest a collapse in microbial-derived immunoregulatory tone. To further contextualize these compositional alterations, we performed KEGG-based predictive metagenomic analyses to infer functional capacities of the microbial communities. In 8-month-old mice, predicted microbial gene modules were enriched for KEGG pathway annotations broadly associated with aging- and immune-related processes, including longevity regulation, RIG-I–like receptor signaling, FoxO signaling, MAPK signaling, and endocannabinoid-associated pathways, relative to 8-week-old controls (Fig. 5F). Importantly, these annotations reflect predicted microbial functional potential rather than direct activation of corresponding host endogenous pathways. Within this framework, lipid metabolism emerged as a particularly disrupted node to regulate Treg stability, with significant alterations in pathways governing unsaturated fatty acid biosynthesis ^36, 38, 39, 40, 41, 42, 43^. In TFAM Het mice, lipid metabolism was profoundly disrupted, with significant alterations observed in pathways governing unsaturated fatty acid biosynthesis, PI3K-Akt signaling, spinocerebellar ataxia pathways, sphingolipid metabolism, primary bile acid biosynthesis, and fatty acid degradation (Fig. 5G). Concurrently, neurotransmission-related pathways, including GABAergic and glutamatergic synapses, were markedly downregulated in aged TFAM Het mice (Fig. 5G). This downregulation aligns with observed behavioral dysfunction and hippocampal atrophy in 8-month-old TFAM Het mice (Fig 3H, 3I, 4A). These findings support a model wherein mitochondrial impairment in regulatory T cells perturbs microbial generation of neuromodulator metabolites, disrupting gut-brain signaling and predisposing the central nervous system to inflammatory remodeling^44, 45, 46, 47^.

To mechanistically link microbial changes with host gene regulation, we performed an integrated correlation analysis between differentially abundant bacterial taxa and host transcriptomes obtained from brain. This analysis uncovered 323 significant microbiota-host gene interactions (FDR-adjusted *p* <0.05), mapping a complex regulatory network that bridges microbial composition and neuroimmune signaling (Fig. 5H). *Helicobacter* species exhibited a strong positive correlation with *Ntsr2*, a neurotensin receptor involved in both neurotransmission and microglial activation, suggesting microbial influence over neural inflammatory circuits. *Lactobacillus* emerged as a central node, negatively correlating with neuroendocrine transcripts (*Pomc*, *Sstr5*, *Ntsr2*) and neuronal developmental regulators (*Elk3*, *Sox4*), while positively correlating with inflammatory mediators (*Sele*, *Mpo*, *Ifitm6*), linking its depletion to neuroimmune destabilization. *Lachnospiraceae_NK4A136_group* was associated with complement-mediated inflammation, showing a positive correlation with *C5ar2* and negative regulation of *Sstr5* and *Elk3*, reinforcing its role in shaping neuroinflammatory tone. To assess downstream inflammatory consequences associated with microbiota-linked alterations in TFAM Het mice, we quantified inflammatory cytokine transcripts in peripheral tissues. IL-1β and TNFR1 expression were significantly elevated in brain tissue (Fig. 5I), indicating the presence of sustained, low-grade neuroinflammatory signaling. These inflammatory changes occurred in concert with marked shifts in microbial community composition and predicted microbial metabolic capacity, including pathways related to lipid and SCFA-associated processes. Together with concurrent shifts in microbial composition and predicted metabolic capacity, these findings support a model in which partial TFAM insufficiency in regulatory T cells destabilizes immune homeostasis and increases susceptibility to age-associated systemic and neuroinflammatory remodeling.

### Modulation of mitochondrial stress pathways attenuates systemic inflammation and aging-associated pathology

To determine whether immune pathology associated with Treg-specific TFAM deficiency is amenable to intervention, we evaluated resveratrol (RSV) treatment in TFAM cKO mice. RSV is a pleiotropic SIRT1-associated compound with established effects on cellular stress responses and inflammatory signaling. Prior studies have demonstrated that the protective effects of RSV across multiple disease contexts are mediated, at least in part, through activation of the PGC-1α–NRF-1–TFAM axis, thereby promoting mitochondrial biogenesis and transcriptional capacity in a SIRT1-dependent manner^48, 49^. Accordingly, RSV was administered intraperitoneally at 7 mg/kg every other day for four weeks (Fig. 6A). Consistent with these reports, RSV treatment was associated with increased transcript levels of *PGC-1α*, *PGC-1β*, and the mitochondrial transcription factors *TFAM* and *TFB2M*, indicating engagement of mitochondrial biogenic programs. At the tissue level, RSV treatment was associated with altered mitochondrial membrane potential, as reflected by MitoSpy staining in skin sections (Fig. 6B), and coincided with a marked reduction in systemic inflammatory burden, evidenced by decreased circulating TNF-α and IL-6 levels (Fig. 6C). In parallel, expression of senescence-associated genes *Cdkn2a* (p16^Ink4a^) and *Cdkn1a* (p21^Cip1^) in splenocytes was reduced, consistent with attenuation of inflammation-linked senescence programs (Fig. 6D-F). Histopathological analysis further revealed diminished inflammatory infiltration and improved preservation of tissue architecture in lungs, liver, and kidney (Fig. 6G).

**Figure 6.**
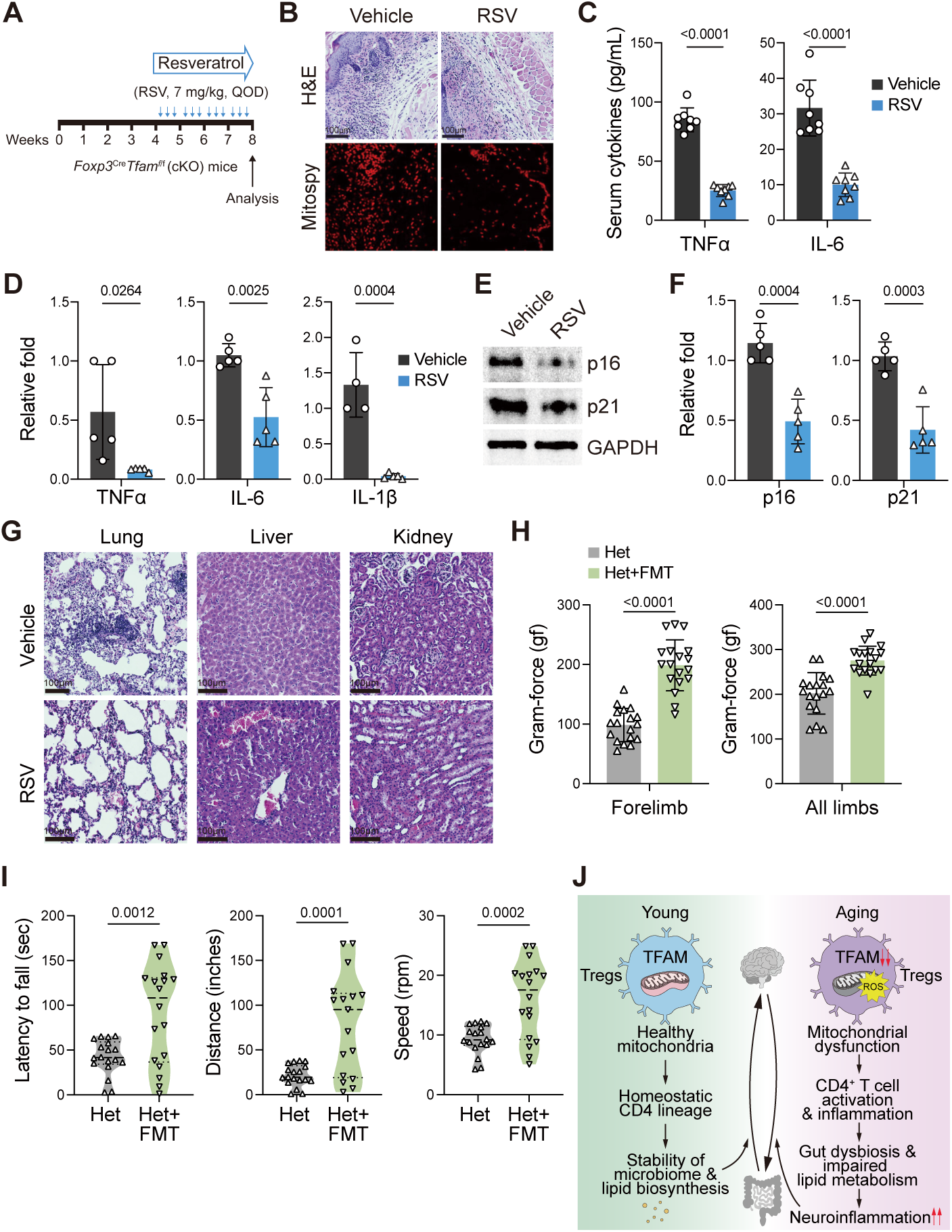
Restoration of mitochondrial fitness mitigates inflammation and functional decline in models of immune aging. (**A**) Schematic of therapeutic intervention in TFAM cKO mice. Mice received intraperitoneal resveratrol (RSV; 7 mg/kg) every other day for 4 weeks. (**B**) Mitospy staining in skin tissue reveals enhanced mitochondrial membrane potential in RSV-treated TFAM cKO mice, indicating improved mitochondrial integrity. (**C**) Serum levels of TNF-α and IL-6 were significantly reduced following RSV treatment, reflecting dampened systemic inflammation. (**D-F**) Quantitative RT-PCR analysis of splenocytes shows reduced expression of senescence markers Cdkn2a (p16^Ink4a^) and Cdkn1a (p21^Cip1^) in RSV-treated mice, consistent with suppression of the senescence-associated secretory phenotype (SASP). (**G**) Histological examination of lung, liver, and kidney shows decreased inflammatory infiltration and preserved tissue structure in RSV-treated animals. (**H**) Experimental design for fecal microbiota transplantation (FMT) in TFAM Het mice. Mice were antibiotic-conditioned for 10 days and subsequently gavage feces from TFAM-sufficient donors over 3 days, followed by a 3-week recovery. (**I**) FMT improved neuromuscular performance, demonstrated by increased rotarod endurance and forelimb grip strength. These improvements were associated with re-expression of Treg-associated genes and suppression of pro-inflammatory programs. (**J**) Working model summarizing the role of TFAM as a central regulator of mitochondrial redox balance, microbiota composition, and immune aging. Restoration of mitochondrial integrity via RSV and microbial reprogramming via FMT ameliorates immunopathology and rejuvenates immune function.

Treg stability is also influenced by microbiota-derived metabolites, especially short-chain fatty acids that support oxidative metabolism and chromatin accessibility at the *Foxp3* locus ^16, 30, 36, 41, 43^. We therefore evaluated whether environmental metabolic inputs show similar corrective trends. Fecal microbiota transplantation (FMT) was performed in TFAM heterozygous (TFAM Het) mice, which exhibit a gradual, non-lethal phenotype more consistent with physiological immune aging than the acute TFAM cKO model. Because regulatory T cell function and peripheral immune tone are strongly influenced by microbiota-derived signals, we reasoned that partial TFAM insufficiency may lower the threshold for microbiota-dependent modulation of host physiology. Antibiotic-conditioned TFAM Het mice received fecal material from TFAM-sufficient donors over three days, followed by a three-week recovery period (Fig. 6H). FMT was associated with improved neuromuscular performance, including increased rotarod endurance and enhanced forelimb grip strength (Fig. 6I). The observed functional improvements indicate that TFAM insufficiency renders host physiology responsive to microbiota-dependent modulation. Together, pharmacologic and microbiota-directed interventions collectively suggest that TFAM-linked mitochondrial competence in Tregs serves as a modifiable nexus linking cellular bioenergetics, immune modulation, and organismal aging. These results align with the perspective that mitochondrial and metabolic support can mitigate the inflammatory and functional aspects of immune aging, rather than effecting a conclusive reversal (Fig. 6J).

## Discussion

The progressive decline in mitochondrial transcriptional integrity is a hallmark of aging, yet its direct impact on immune regulation and cellular fate remains insufficiently characterized^50, 51, 52^ ^17, 32, 53^. Specifically, whether the decreasing mitochondrial transcriptional capacity within Tregs directly initiates immune aging, or merely reflects broader senescent cues remain unresolved. Here, we mechanistically establish TFAM as a central molecular rheostat of Treg identity and systemic immune homeostasis. Using Treg-specific physiologically relevant graded TFAM insufficiency, we demonstrate that progressive disruption of mitochondrial transcriptional output rewires CD4⁺ T cell fate, reshapes the gut microbial ecosystem, and primes the central nervous system for neuroinflammatory remodeling.

A major limitation in immune aging research has been the absence of physiologically relevant models that can selectively dissect the cell-intrinsic consequences of mitochondrial dysfunction within Tregs across the aging trajectory. Existing models involving global TFAM deletion or tissue-specific knockouts in high-turnover compartments often result in rapid lethality, systemic cytokine storm, and developmental collapse, precluding longitudinal assessment of immune remodeling^12, 16, 24, 54, 55^. Similarly, Treg ablation models such as Foxp3^DTR^ or DEREG mice induce severe autoimmunity and fail to capture the progressive, transcriptionally encoded dysfunction of suppressive Treg identity observed during physiological aging^19, 56^. Other aging models, including Ercc1^-/Δ^, BubR1^H/H^, and Zmpste24^-/-^, reflect accelerated DNA repair defects or nuclear envelope instability, but do not reproduce the slow, cell-specific decline in mitochondrial transcriptional output characteristic of aging Tregs^57, 58,59^.

To overcome these constraints, we developed the Foxp3^Cre^Tfam^f/+^ mouse, which induces a stable ∼50% reduction in TFAM within Foxp3⁺ Tregs, consistent with the age-related TFAM decline reported in human PBMCs. Unlike Foxp3^Cre^Tfam^f/f^ models that succumb to early mortality, TFAMHet mice remain viable, thereby enabling longitudinal tracking of mitochondrial insufficiency and its systemic consequences. With age, TFAMHet mice exhibit canonical features of organismal aging kyphosis, sarcopenia, reproductive decline, and neuromuscular impairment alongside immune aging hallmarks, including elevated p16^Ink4a^, p21^Cip1^, increased circulating IL-6 and TNF-α, and a shift of the CD4⁺ T cell compartment toward CD44^hi^CD62L^lo^ effector memory phenotypes. In the CNS, TFAMHet mice show heightened microglial activation (CX3CR1⁺MHC-I^hi^) and hippocampal neuronal loss, consistent with peripheral-to-central immunometabolic disruption (Fig. 3-4). Collectively, this model provides a genetically precise, temporally scalable system to dissect how mitochondrial transcriptional fidelity in Tregs orchestrates immunological aging, gut–brain communication, and tissue integrity. The TFAMHet mouse thus offers a tractable platform for probing mitochondria-driven immunoregulation with translational relevance to human aging.

While mitochondrial metabolism is known to sustain FOXP3 expression and Treg function, the direct role of mitochondrial genome integrity specifically TFAM-dependent transcriptional control, in orchestrating systemic CD4⁺ T cell fate has remained undefined. As aging coincides with both mitochondrial erosion and effector CD4⁺ expansion, the mechanistic bridge linking these processes has not been elucidated. By leveraging single-cell RNA sequencing spleens from TFAMcKO mice, we uncover a profound reshaping of the CD4⁺ T cell landscape. TFAM loss in Tregs drives expansion of cytotoxic Gzmb⁺Ccl5⁺ subsets, S100a6⁺ effector memory (TEM) cells, and Icos⁺Rgs1⁺ aged-like Tregs (aTregs), coupled with depletion of early-stage Tcf7⁺Lef1⁺ naïve and Tnfsf8⁺ activated/exhaustion-associated populations. These alterations are consistent with transcriptionally encoded, lineage-intrinsic reprogramming rather than being explained solely by a secondary inflammatory milieu, as revealed by pseudotime reconstruction and RNA velocity mapping. Notably, TFAM deficiency enforces commitment along an alternative P2 trajectory enriched for neuroinflammatory and Th17-like gene modules, including *Anxa2*, *S100a4*, and *Ccl5*, while repressing transcriptional brakes such as *Satb1* and *Bcl2*, suggesting a premature exit from naïve states and loss of regulatory restraint. Mechanistic interrogation through gene regulatory network (GRN) analysis reveals a collapse of Foxp3-lineage stability in TFAM-deficient Tregs, marked by activation of inflammatory regulons (*Nfkb2*, *Nfatc1*, *Runx2*), cytolytic transcriptional drivers (*Runx3*, *Prdm1*), and stress-response regulators (*Ddit3*, *Egr1*, *Junb*). The induction of Ddit3, a hallmark of mitochondrial unfolded protein response (UPR^mt^), together with RIPK1/RIPK3/MLKL phosphorylation and caspase activation, is consistent with activation of mitochondrial stress linked cell-death programs (including necroptosis and apoptotic components) that could propagate paracrine stress to surrounding CD4+ subsets. This remodeling is accompanied by transcriptional enrichment for neurodegenerative signatures, including PI3K-Akt, FoxO, and oxidative phosphorylation pathways supporting a molecular link between mitochondrial erosion in Tregs and CNS-relevant inflammatory programming. Our findings delineate TFAM-dependent mitochondrial transcription as a master regulator of CD4⁺ T cell fate diversification, acting upstream of immunosenescence and neuroinflammation. By defining mitochondrial instability in Tregs as a transcriptional checkpoint that governs effector reprogramming, our study uncovers a novel axis of immune aging in which fidelity of Treg mitochondrial transcription actively restrains the emergence of pro-inflammatory and potentially neurotoxic CD4⁺ states.

A central unresolved question in immunometabolism and aging is whether mitochondrial dysfunction in Tregs actively shapes gut microbiota composition and influences the gut–brain axis, or whether microbiome alterations are merely downstream byproducts of systemic aging^35, 36, 37^. While prior studies have established that SCFAs, particularly butyrate and propionate, produced by gut microbiota promote FOXP3 expression and enhance Treg function through epigenetic remodeling, the reciprocal influence of Treg mitochondrial state on microbial ecology has remained mechanistically undefined^36, 41, 42, 43, 46^. Our study addresses this critical gap by demonstrating that partial loss of mitochondrial transcriptional control in Tregs via TFAM haploinsufficiency is sufficient to disrupt microbial-metabolite circuits essential for immune regulation and neuroimmune integrity. In TFAMHet mice, we observed a marked depletion of SCFA-producing bacterial taxa, including *Lactobacillus murinus* and *Odoribacter*, which are essential for maintaining Treg epigenetic programming and FOXP3 stability^36, 44,60^. This dysbiosis coincided with broad remodeling of gut lipid metabolic pathways, particularly involving bile acid biosynthesis, sphingolipid metabolism, and signaling cascades such as PI3K-Akt and FoxO, that are frequently implicated in aging-related immune and neuronal dysfunction^61, 62^.

Importantly, our integrative analysis combining single-cell RNA-seq, lipidomics, microbiome profiling, and host–microbiota transcriptome correlation revealed that Treg-intrinsic TFAM insufficiency is associated with a systems-level disruption of gut–brain communication. Neurotransmission pathways, especially GABAergic and glutamatergic circuits critical for CNS function, were significantly disrupted in TFAMHet mice, aligning with hippocampal atrophy and behavioral decline. Furthermore, correlation mapping uncovered 323 robust associations between microbial taxa and neuroimmune gene networks. Notably, *Helicobacter* spp. positively correlated with *Ntsr2*, a neurotensin receptor linked to microglial activation, while *Lactobacillus* abundance negatively correlated with *Pomc*, *Sstr5*, *Ntsr2*, and transcriptional regulators *Elk3* and *Sox4*, underscoring a microbial influence on CNS signaling cascades^41, 63^. Elevated expression of inflammatory mediators such as *Il1β* and *Tnfr1* in TFAMHet tissues further support the emergence of a chronic, low-grade inflammatory state driven by Treg mitochondrial insufficiency. Recent studies have shown gut–brain axis is a complex, bidirectional communication network that links the gastrointestinal tract and the central nervous system through neural, hormonal, and immune pathways. Together, these findings support a previously underappreciated framework linking Treg mitochondrial integrity with lipid-associated metabolic programs and microbial community structure. In this model, Treg-specific TFAM insufficiency is associated with coordinated changes in microbial composition, reduced representation of SCFA-producing taxa, and heightened inflammatory signaling, reflecting a destabilized immunometabolic state. Rather than defining a closed causal feedback circuit or a resolved gut-brain immune equilibrium, our data position mitochondrial fidelity in Tregs as a critical stabilizing influence on immune–microbial interactions that shapes susceptibility to systemic and neuroinflammatory remodeling during aging.

Finally, we provide a proof-of-concept that mitochondrial dysfunction in Tregs is therapeutically actionable through redox restoration and microbiome modulation. In the TFAMcKO model, resveratrol an established activator of SIRT1 and PGC-1α signaling enhanced mitochondrial biogenesis, restored mitochondrial membrane potential, reduced senescence markers, and suppressed systemic inflammation. These findings align with prior reports demonstrating resveratrol’s capacity to improve mitochondrial function and mitigate age-associated inflammation across tissues^49, 64^. In the chronic TFAM Het model, fecal microbiota transplantation (FMT) restored SCFA-producing taxa, improved neuromuscular performance, and normalized immune signatures. While FMT is increasingly recognized for its ability to modulate systemic immunity and neurocognitive health via the gut-brain axis^65, 66^, its role in regulating Treg mitochondrial function remains poorly defined. Our data suggests that microbiota-derived metabolites may enhance peripheral tolerance or indirectly support Treg mitochondrial homeostasis. Together, these findings position Treg-intrinsic mitochondrial decline as a reversible driver of inflammaging and support the translational potential of combining metabolic and microbial interventions to combat age-related immune dysfunction (Fig. 6).

In sum, our study redefines Treg mitochondrial integrity as a central axis of immunological aging. By uncovering TFAM as a lineage-specific molecular rheostat that governs immune suppression, microbial composition, and neuroimmune resilience, we provide a unifying framework for the mitochondrial regulation of systemic aging. These findings open translational opportunities for mitochondria-targeted immunotherapies in chronic inflammatory diseases, neurodegeneration, and age-associated immune decline.

## Methods

### Animals

Foxp3^YFPCre^ (Stock No: 016959) and TFAM^f/f^ (Stock No: 026123) mice were obtained from Jackson Laboratory (Bar Harbor, ME) and bred at the UND animal facility to create Foxp3^Cre^TFAM^f/f^. All mice used in this study were from C57BL/6 backgrounds. Animals were kept and bred in the UND Animal facility in a pathogen-free unit with a 12-hour light cycle, temperature, and humidity control. Efforts are being made to reduce animal suffering and needless usage, in accordance with the Institutional Animal Care and Use Committee’s (IACUC) and National Institutes of Health’s Guide for the Care and Use of Laboratory Animals standards.

### Histology, immunohistochemistry, and acquisition of images

Formalin-fixed, paraffin-embedded slices of various tissues were deparaffinized on slides using xylene twice for 5 minutes each. Following that, they were switched to 100% ethanol twice for 3 minutes each and then to 95%, 75%, 65%, 50%, 35%, and 25% ethanol for 3 minutes each. Rinse the slides for ten minutes under running tap water. The excess water was removed using a wiper (KIMTECH SCIENCE) and 10% fetal bovine serum (FBS) in PBS was applied to the sections of the slides and incubated for 30 minutes in the dark at room temperature (RT). FBS was gently drained from the slides and cleaned with PBS (1X). MitoSpy (excitation (ex)/emission (em) at 577nm/598nm, MitoSpyTM Red CMXros (Cat#424801, BioLegend), After incubation, slides were washed twice with 1X PBS and a single drop of mounting media (Fluoromount-G^TM^ with DAPI, Cat# 00-4958-02, Thermo Fisher Scientific) was applied to the tissue sections and the coverslip was dipped into the mounting solution. For microglia (IBA1⁺) and neuronal (NeuN⁺) immunostaining, mouse brains were fixed in 4% paraformaldehyde (PFA) in 1× PBS, cryoprotected in 30% sucrose, and cryo-sectioned. Tissue sections were blocked in 0.2% Triton X-100 and 5% normal goat serum in 1× PBS for 1 hour at room temperature, followed by incubation with primary antibodies rabbit monoclonal anti-IBA1 (Cat# 17198, 1:1000) and mouse monoclonal anti-Neuronal N (Cat# 94403, 1:1000) diluted in 0.2% Triton X-100 and 3% normal goat serum in PBS for 24 hours at 4°C. After three 20-minute washes in PBS, sections were incubated with appropriate secondary antibodies anti-Rabbit CF^™^594 antibody (Cat# SAB4600107) and anti-mouse CF^™^488 antibody (Cat# SAB4600388) for 1 hour at room temperature. Nuclei were counterstained using a single drop of Fluoromount-G™ with DAPI (Cat# 00-4958-02, Thermo Fisher Scientific) and sealed with nail polish. Fluorescence images were acquired using a Leica Thunder Imaging System and an Olympus TIF fluorescence microscope. Whole-slide histological scans were performed using a Hamamatsu NanoZoomer at the UND Histology Core. Quantification of immunofluorescence signals was carried out using NIH ImageJ software.

### Western blot

Proteins were isolated from mouse tissue and/or single cells and homogenized in RIPA lysis buffer containing a 1% protease inhibitor cocktail^67, 68, 69^. The protein concentration was determined according to the manufacturer’s instructions and our previously published protocol using the Bio-Rad protein assay reagent. SDS-PAGE was used to extract a set quantity of cellular protein and electroblotted onto an Immobilon-P transfer membrane (Cat# IPVH00010). The immunological blot was blocked for 1 hour at room temperature with 5% nonfat dry milk in TBST (25 mM Tris-HCL, pH 7.4, 1.5 M NaCl, 0.05% Tween-20), followed by overnight incubation with primary antibodies at 4 °C. The next day, the membranes were washed five times at six-minute intervals and incubated for one hour with HRP-conjugated secondary antibody (A16172, Invitrogen) (1:1000) in 3% nonfat dry milk in TBST. The blots were then washed five further times with TBST and developed with enhanced chemiluminescence. The following primary antibodies were bought from Cell Signaling Technologies (Cell Signaling Technology, MA 01923, USA): RIP (D94C12) Rabbit mAb #3493, MLKL (D2I6N) Rabbit monoclonal antibody #14993, RIPK3 (E1Z1D) Rabbit monoclonal antibody #13526, p-RIP (D1L3S) Rabbit monoclonal antibody #65746, p-MLKL Rabbit monoclonal antibody (D6H3V) #91689, p-RIP3(D6W2T) Rabbit monoclonal antibody #93654, p21 Waf1/Cip1 (12D1) rabbit monoclonal antibody #37543, p53 (1C12) Mouse monoclonal antibody #2524, CDK4 (D9G3E) Rabbit monoclonal antibody #12790, PCNA (D3H8P) XP® Rabbit monoclonal antibody #13110, p16 INK4A (D3W8G) GAPDH (D16H11) XP® Rabbit mAb #5174.

### RNA isolation and qPCR

Total RNA was isolated from mouse tissues or single cells. The tissues were minced with a scissor and added to 500 µL of TRIzol reagent (Ambion, cat# 15596018, Life Technologies, USA)^67, 68, 69^. A Fisher Scientific Sonic Dismembrator 100 was used to fully homogenize tissue solutions (American Laboratory Trading, East Lyme, CT 06333. USA). The upper layer was removed, and the RNA was precipitated with an equivalent volume of isopropanol and incubated at 55°C for 10 minutes before being resuspended in 50 µL of RNase-free water (Promega, Madison, WI, USA). We determined the concentrations of RNA using a Smart-Spec plus spectrophotometer (Bio-Rad). Then, using a High-Capacity cDNA Reverse Transcription Kit (Cat#: 4368814, Applied Biosystem, Thermo Fisher Scientific), cDNA was generated from 0.5-1 µg of total RNA in a total volume of 25 µL. The RT-PCR program was run for 5 minutes at 25 °C, 30 minutes at 42 °C, and 30 minutes at 95 °C. The cDNA templates were further diluted in water two-three times. For qPCR, two microliters of diluted templates were utilized. A 96-well PCR plate (Cat#: MLL9601, Bio-Rad) was used for the qPCR. All genes were amplified twice, and each sample was analyzed in triplicate using 2X SYBR Green PCR mix (Cat#: B21202, https://Bimake.com) (Supplementary Table 6). After normalization to *Gapdh*, the CT value was utilized to compute the fold change in RNA abundance. qPCR reactions were performed using the Aria Mx Real-Time PCR equipment (Agilent).

### Flow cytometry and cytokine analysis

Single-cell suspensions were prepared from mouse spleen. To assess CD4^+^ T cell activation, we stained single-cell suspensions cell surface staining in 50µL FACS buffer (1% BSA, 2 mM EDTA, and 0.1% sodium azide in PBS, pH 7.4) (5-10x10^5^ cells/tube) and incubated with antibodies for 30 minutes at 37°C. (106 cells/50 mL) with anti-CD45 PercpCy5.5 (Cat# 103132, Clone#30F11, BioLegend), anti-CD4 PE (Cat#100408, Clone#GK3.5, BioLegend), anti-CD44 Alexflour700 (Cat#103026,IM7, BioLegend), and anti-CD62L PeCy7 (Cat#104418, Clone# MEL/4, BioLegend) on ice for 30 minutes (SupplementaryTable 6). Stained cells were then washed twice with 500µL of ice-cold FACS buffer to eliminate binding antibodies before being analyzed immediately using a BD FACSymphony A3Flow Cytometer. Annexin V APC staining cells were incubated with Annexin V APC (Cat#640920, BioLegend) for 3 minutes in the ice-cold FACS buffer and then washed. Followed by Propidium Iodide (Cat#6421301, BioLegend) added right before analyzing the cells by FACS. MitoSpy (Cat# 424801, BioLegend) staining was carried out in accordance with the manufacturer’s recommendations. Cells were resuspended in MitoSpy in 300-400 µL of warm 1X PBS and incubated at 37 °C for 30 minutes. Cells were then washed twice with 1X PBS before being analyzed using a BD FACSymphony A3 Flow Cytometer on stained and unstained samples (BD Biosciences). For inflammatory cytokines detection, sera were collected from mice blood and measured by the LEGENDplex mouse inflammation panel according to the manufacturer’s instructions (Cat#740446, BioLegend). Samples were read on the BD FACSymphony Flow Cytometer.

### *In vitro* Treg suppression assay

Tregs (CD4^+^CD25^+^) cells were isolated from the spleen of sex-matched TFAM cKO and TFAM^f/f^ (control) mice and Tconv (CD4^+^CD25^-^) from TFAM control mice using BD FACS ARIA^TM^llu at the UND Flow Cytometry Core: anti-CD4 PE (Cat#100408, Clone#GK3.5, BioLegend), anti-CD25 (Cat#102030, clone PC61, BioLegend). Afterward, 5x10^4^ Tconv (CD4^+^CD25^-^) cells from TFAM control were labeled with CFSE Cell Division Tracker Kit (Cat# 423801, BioLegend) and cultured with mitomycin-treated wild type splenocytes. To detect Treg suppressive effect on these labeled Tconv (CD4^+^CD25^-^) cells culture, we added Tregs (CD4^+^CD25^+^) from the TFAM cKO and TFAM^f/f^ (control) for 60 hours at 37°C with 2.5 mg/mL Dynabeads^TM^ Mouse T-Activator CD3/CD28 for T-Cell Expansion and Activation bead (Cat# 1145). After 60 hours, the CD4 cell proliferation was observed by FACS. Suppressive capacity was calculated based on the relative proportion of proliferating Tconv cells compared with cultures lacking Tregs and is presented as percent suppression, as our previously described study^69^.

### Cell isolation and sequencing library construction using the 10x Genomics Chromium platform

Cells were isolated from spleens from WT and cKO mice by grinding with 70 µm cell strainers (Cat#22363548, Fisher brand). ACK RBC lysis buffer (Cat#A10492-01, Gibco) was used to lyse RBC. Trypan blue (Cat#25900Cl, Corning) staining confirmed cell viability. The 10X Genomics Chromium system was used to create 3’ single-cell gene expression libraries (v3.0) from WT and cKO spleen single-cell suspensions. On an Illumina HiSeq X Ten, pair-end 150 bp sequencing was used to generate high-quantity data (Illumina, San Diego, California, USA). Then, using Cell Ranger software v6.0.0, the output Illumina base call files from sequencing runs were combined and converted to FASTQ files using Illumina’s bcl2fastq.

### Single-cell RNA-seq data alignment and sample aggregating

RNA reads from the 10X Chromium platform with a sequence quality of less than 30 were filtered out and then mapped to the mm10 mouse reference genome using the CellRanger software (v6.0.0). Seurat (v4.0.2) was used to read the individual sample output files from CellRanger Count^70^. Cells were further filtered out of the downstream analysis using the following criteria: *nFeature RNA <200 & nFeature RNA >4000 & percent.mt >25*. Scrublet (v0.2.3)^71^ was used to detect probable doublets and then *CellCycleScoring in Seurat was used* to define the cell cycle phase by using a published list of cell cycle genes^72^. *SCTransform* (v0.3.2) from Seurat was used to independently normalize the gene-cell count matrices with default parameters^73^. Dimension reductions, including principal component analysis (PCA), clustering, and t-distributed Stochastic Neighbor Embedding (t-SNE) projections (resolution =0.7) were then performed on the normalized data. By following a similar pipeline, subclusters for CD4^+^ T cells were identified.

### Cell type annotation and differential expression analysis

To determine the cell type of each cluster, the *FindAllMarkers* function in Seurat was used to find the cluster’s gene markers. Cell types were allocated manually after consulting the CellMarker^74^ and PanglaoDB^75^ databases. MAST (v1.18.0)^76^ was utilized to conduct differential analysis on each cell type to find the genes that were differentially expressed between WT and cKO cells with the adjusted p-value <0.05 and absolute log2FC >0.25. The Gene Set Enrichment Analysis (GSEA) with the Kyoto Encyclopedia of Genes and Genomes (KEGG) pathway annotation was done using the richR package (https://github.com/hurlab/richR), with a p-value of 0.01 used as the cutoff value to identify significant KEGG pathways.

### Single-cell trajectory reconstruction for CD4^+^ T cell

The Monocle2 R package (v2.20.0, http://cole-trapnell-lab.github.io/monocle-release/) was used to reconstruct cell fate choices and pseudotime trajectories. The gene-cell matrix of UMI counts was sent to Monocle, which then created an object using its new Cell Data Set function (*lower Detection Limit = 0.5, expression Family = nonbinomial.size*). We evaluated the size factors and dispersion of gene expression. We next used the differentialGeneTest module to conduct differential expression analysis in order to determine the ordering of genes using the parameter “*fullModelFormulaStr = ∼celltype*,” and then maintained genes with q values less than 1E-3 for downstream analysis. Following the *setOrderingFilter* function, the *reduceDimension* module with the argument “*method = DDRTree*” was used to reduce the dimensionality of the data. When *orderCells* is executed, the root state variable may be adjusted to reflect the known biological background. Following that, the *plot_cell_trajectory* and *plot_pseudotime_heatmap* functions were used to visualize the data further.

### RNA velocity analysis

Velocyto (v0.17.17) was used to analyze the velocity of RNA in cells^77^. This method utilizes the relative amount of unspliced and spliced mRNA as a predictor of future cell state. Annotation of spliced and unspliced reads was conducted using the Python script velocyto.py in the Cell Ranger output folder utilizing the Velocyto pipeline. Then, the scvelo (v0.2.3)^78^ was used to determine the CD4^+^ T cell velocity. The previously normalized data served as the basis for calculating first and then order moments for each cell in relation to its closest neighbors (*scvelo.pp.moments(n pcs = 30, n neighbors = 30)*). Following that, the velocities were calculated, and the velocity graph was produced using the *scvelo.tl.velocity* and *scvelo.tl.velocity* graph functions with the mode set to ‘stochastic’. Finally, using the *scvelo.tl.velocity* embedding function, the velocity field was projected onto the existing t-SNE coordinates.

### Transcription factor regulons prediction

Gene regulatory network (GRN) was generated using pySCENIC package (v0.11.0)^79^. Briefly, the raw expression matrix for the cells of all samples was filtered by keeping genes with a total expression greater than 3*0.01*(number of cells). Then the *pyscenic.grn* command was used with *GRNboost2* method and default options and a fixed seed to derive co-expression modules between transcription factors and potential targets. The following database: cisTarget databases (*mm10 refseq-r80 10kb_up_and_down_tss.mc9nr.feather, mm10 refseq-r80 500bp_up_and_100bp_down_tss.mc9nr.feather*), the transcription factor motif annotation database (*motifs-v9-nr.mgi-m0.001-o0.0.tbl*) and the list of mouse transcription factors (*mm_mgi_tfs.txt*) were downloaded for the analysis. A regulon is a group of target genes regulated by a common transcription factor. AUCell (Area Under the Curve) scores (regulon activities) in each cell were computed with *pycenic.aucell* command (default parameters). These scores were then binarized into either “on” or “off” states, by setting an optimized threshold on the distribution of each regulon among all cells using the *skimage.filters.threshold_minimum* function.

### Bulk RNA-seq data processing

Low quality (Q<30) reads, and sequencing adapters were removed with Trimmomatic software v0.36^80^ from raw reads. Clean reads were then mapped to the mouse reference genome (mm10) by using HISAT2^81^. FeatureCounts^82^ were used to summarize the unique mapped reads to mouse genes. Differential gene expression analysis was performed using DESeq2^83^ with adjusted p-value <0.05 and fold change >2 as the cutoff for identifying differentially expressed genes (DEGs). Gene Set Enrichment Analysis (GSEA) on KEGG pathway data was performed by using the richR package and the adjusted p-value <0.05 was chosen as the cutoff value to select significant KEGG pathways.

### Microbiome analysis

Microbial DNA samples were collected as our vius published protocol^33^. Samples were sequenced using Oxford Nanopore Technology. Raw sequencing data underwent quality control using Porechop (v1.0.0) to remove adapter sequences, followed by length filtering with NanoFilt to retain sequences between 1,200 and 1,800 base pairs. Quality metrics were assessed using NanoStat. Chimeric reads were identified and removed through an all-versus-all alignment approach using minimap2 (v2.17) with Oxford Nanopore-specific parameters (-x ava-ont -g 500), followed by filtering with yacrd (coverage threshold 4, noise threshold 0.4). Cleaned reads were mapped against a reference database using minimap2 (-x map-ont), and high-confidence alignments were retained using custom filtering scripts from the Spaghetti pipeline. OTU tables and taxonomy assignments were generated for downstream analysis.

Microbiome analysis was performed in R (version ≥4.0.0) using the phyloseq package. Alpha diversity (Observed richness, Simpson, and Shannon indices) was calculated at the genus level. Beta diversity was assessed using Bray-Curtis dissimilarity and visualized with Principal Coordinates Analysis (PCoA). Differential abundance analysis between experimental groups (P8M, P8W, PT, and PY) was conducted using ALDEx2 with 128 Monte Carlo instances. To identify significantly differentially abundant bacterial taxa between groups, we used the ALDEx2 package^84^. This approach was chosen to properly handle the compositional nature of microbiome data. Four comparisons were performed: P8M vs P8W, PT vs PY, P8W vs PY, and PT vs P8M. For each comparison, Monte Carlo sampling with 128 Dirichlet instances was used to account for technical variation (mc.samples = 128), with all features included in the geometric mean calculation (denom = “all”).Bacterial taxa were considered significantly differentially abundant if they met both thresholds: p-value < 0.05 (Welch’s t-test) and effect size > 0.8. The effect size threshold represents cases where the between-group difference exceeds 80% of the maximum within-group dispersion^85^. Pathway functionality was predicted from the taxonomic data using PICRUSt2 (Phylogenetic Investigation of Communities by Reconstruction of Unobserved States), allowing for inference of KEGG pathways present in the microbial communities based on marker gene sequencing data. Significant pathways (p <0.05) were visualized using heatmaps.

### Antibiotic Treatment and Fecal Microbiota Transplantation (FMT)

To deplete endogenous microbiota, mice were administered a broad-spectrum antibiotic cocktail in drinking water for two weeks: ampicillin (10 mg/mL, A1593, MilliporeSigma), metronidazole (10 mg/mL, PHR1052, MilliporeSigma), and vancomycin hydrochloride (5 mg/mL, SBR00001, MilliporeSigma). Fecal material for transplantation was collected from donor mice aged 8 weeks (young) or 2 years (aged), snap-frozen, and stored at -80°C. Following antibiotic treatment, recipient C57BL/6 mice were housed in sterile, autoclaved cages and orally gavaged with 300–400 μL of freshly prepared fecal suspension from untreated donor mice. The suspension was generated by homogenizing fecal pellets in sterile water. Control groups received either sterile water or autologous fecal homogenates.

### Grip strength analysis

Neuromuscular strength was assessed using the grip strength test. The forelimb and hindlimb grip strengths of mice were assessed by using a grip strength meter from San Diego Instruments. Each mouse was tested in three trials, and the force was recorded as grams-force (gf, 1 gf = 9.8 x 10^-3^ Newtons).

### Open field test

Locomotor activity was measured in the open field test. A mouse was placed in the empty square cage from San Diego Instruments. Total distance (m) covered during a 10-min period was recorded and calculated by ANY-maze software (version 7.1).

## Data availability

Datasets generated in this study by RNA-seq and scRNA-seq are accessible at GSE203143 (Reviewer access token: ozebasggjbyrpsf) and GSE197973 (Reviewer access token: clajaakoztkhtyh), respectively. All data associated with this study are present in the paper or the Supplementary Materials.

## Code availability

All data analysis scripts are available on the GitHub (https://github.com/guokai8/TFAM).

---

**Supplementary Fig. 1:**
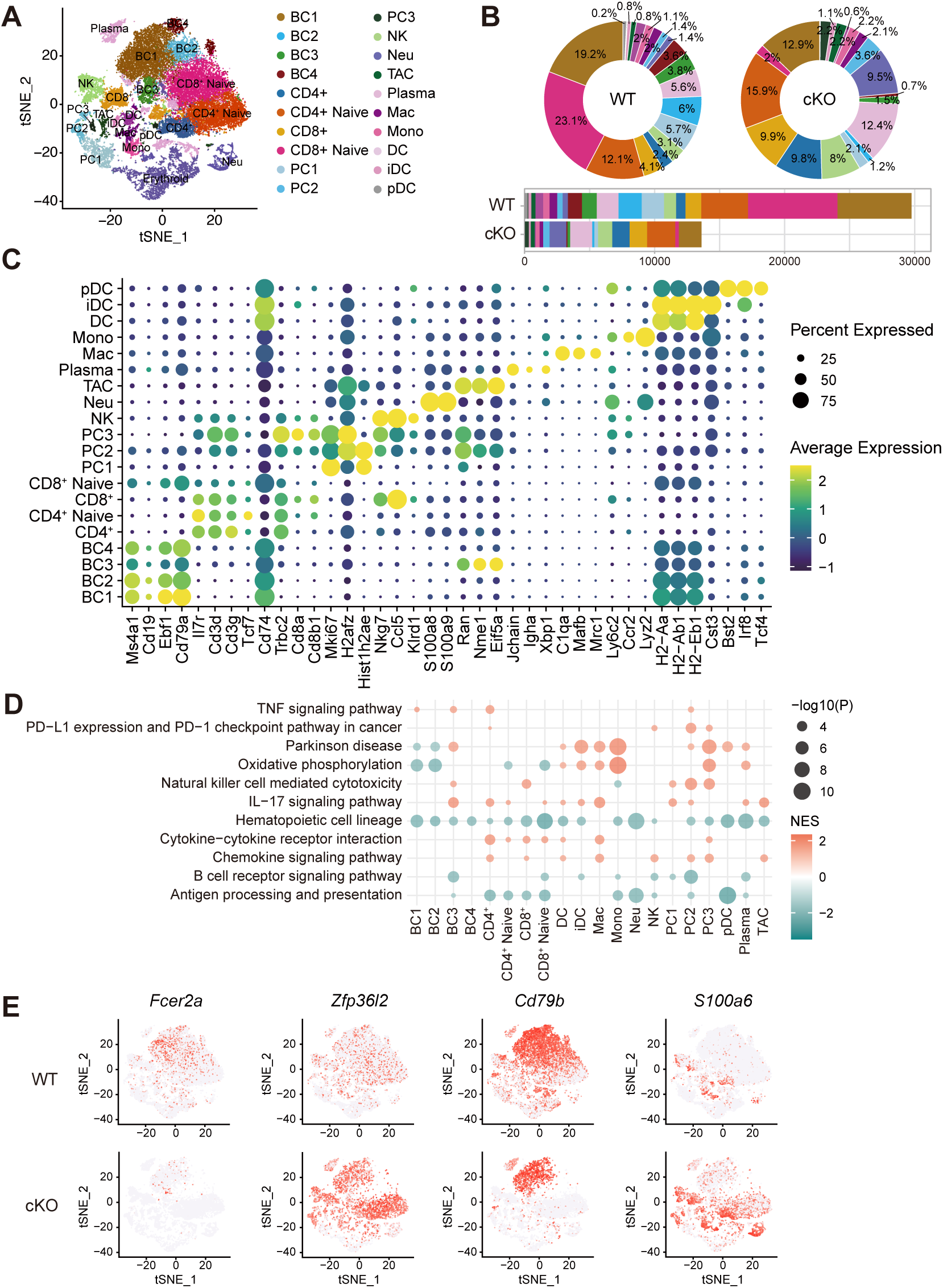
Landscape of scRNA-seq analysis of WT and cKO spleen. **A.** t-SNE projections of single-cell transcriptomes from WT and cKO mice before Erythroid cells were removed, annotated by cell type. Each dot denotes a distinct cell. **B.** Cell subset proportions (top) and numbers (bottom) for WT and cKO spleens. **C.** Dot plot of mean expression of canonical marker genes for 20 major lineages from tissues of each origin, as indicated. **D.** Gene Set Enrichment Analysis (GSEA) of KEGG pathways for all cell types between cKO and WT (Only age-related pathways are selected). Color stands for the up-regulated (red) or down-regulated (blue) in cKO spleen. Enriched terms were identified as significant at a p-value ≤ 0.01. **E.** Expression of aged-related genes (*Fcer2a*, *Cd79b*, *Zfp36l2*, and *S100a6*) on Figure 1B t-SNE embedding. Each dot corresponds to a single cell.

**Supplementary Fig. 2:**
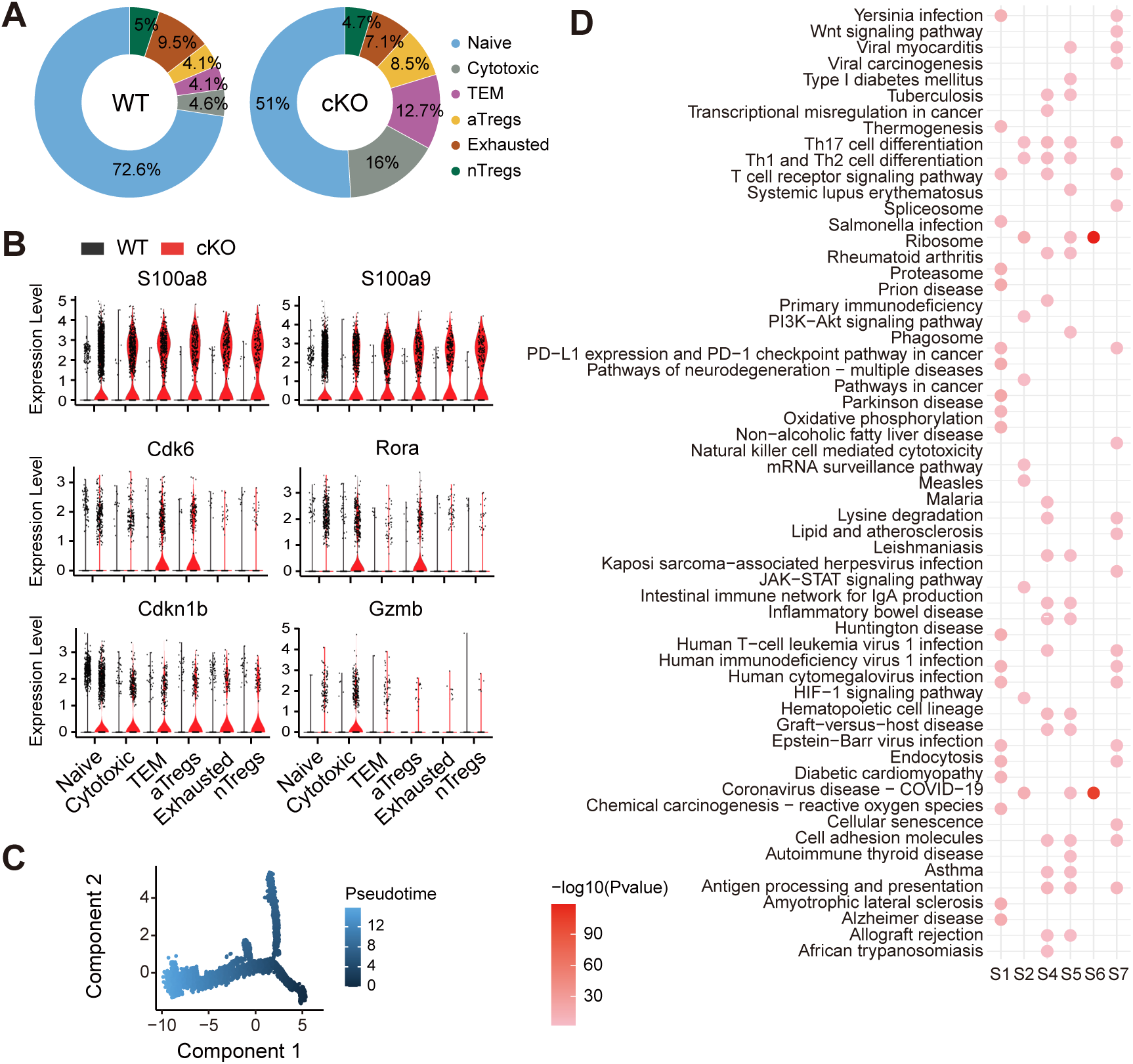
scRNA-seq results show the cellular heterogeneity in CD4^+^ T cells. **A.** Relative proportion of CD4^+^ T cells among WT and cKO spleen, color stands for cell type. **B.** Violin plot showing the expression profile of age-related genes among all CD4^+^ T cells. **C.** Monocle 2 prediction of CD4^+^ T cell subsets developmental trajectory with pseudotime. Each dot represents an individual cell. **D.** Functional enrichment analysis of KEGG pathways with up-regulated differential expressed genes for the cell states. Only the top 20 of the enriched pathways were plotted from each state.

**Fig. 3:**
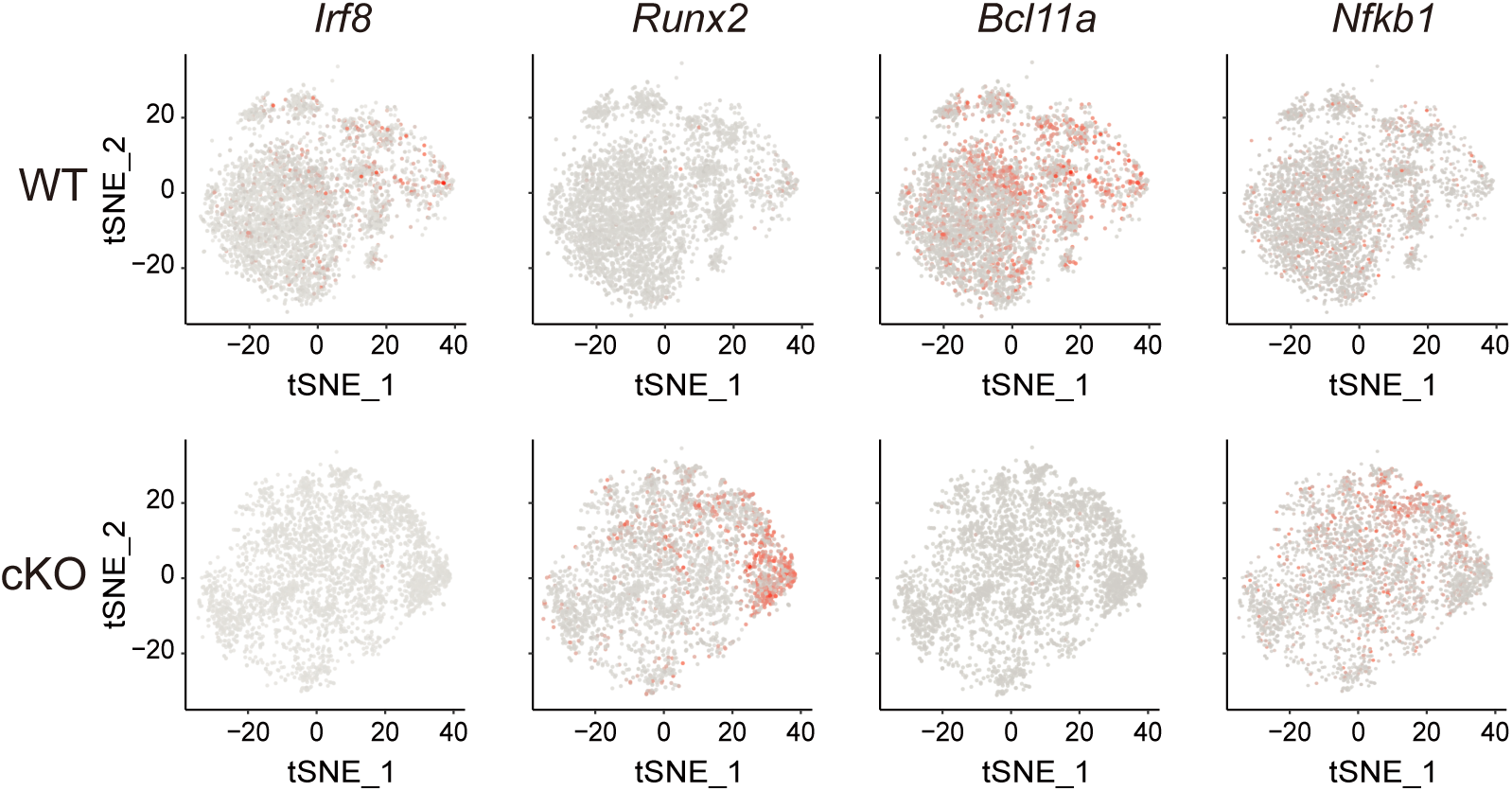
t-SNE projection of binary regulon activity. t-SNE plots visualize the binarized regulon activity states of Irf8, Runx2, Bcl11a, and Nfkb1 across single cells, highlighting distinct transcriptional regulatory modules and cell-type-specific activation patterns.

**Supplementary Fig. 4:**
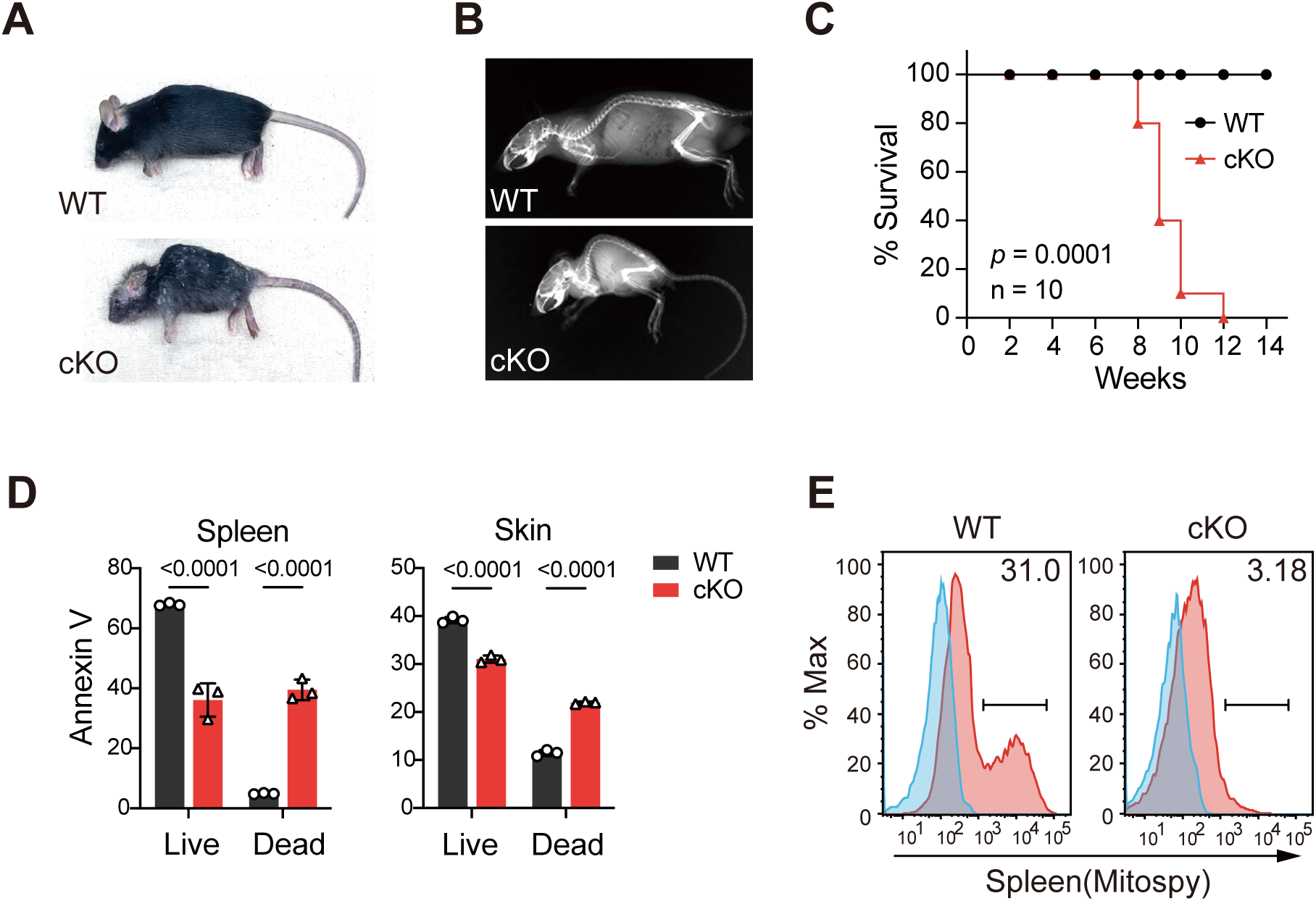
TFAM deletion in Foxp3⁺ Tregs induces systemic pathology and accelerates immune aging in Foxp3^Cre^Tfam^f/f^ mice. **A.** Representative photograph showing the deteriorated physical appearance of a cKO mouse (right) compared to an age- and sex-matched WT littermate (left) at 8 weeks of age. **B.** X-ray image depicting increased spinal curvature in the cKO mouse (right) relative to a WT littermate (left) of the same age and sex. **C.** Kaplan-Meier survival analysis comparing WT (n=10; M/F) and cKO (n=10; M/F) mice, revealing reduced lifespan in cKO animals. **E.** Flow cytometric analysis of Annexin V staining in spleen and skin cells from WT and cKO mice, indicating increased cell death in cKO tissues. **F**. Mitospy staining of splenocytes from WT and cKO mice, assessed by flow cytometry and presented as a fluorescence intensity histogram, showing mitochondrial alterations in cKO cells.

**Supplementary Fig. 5:**
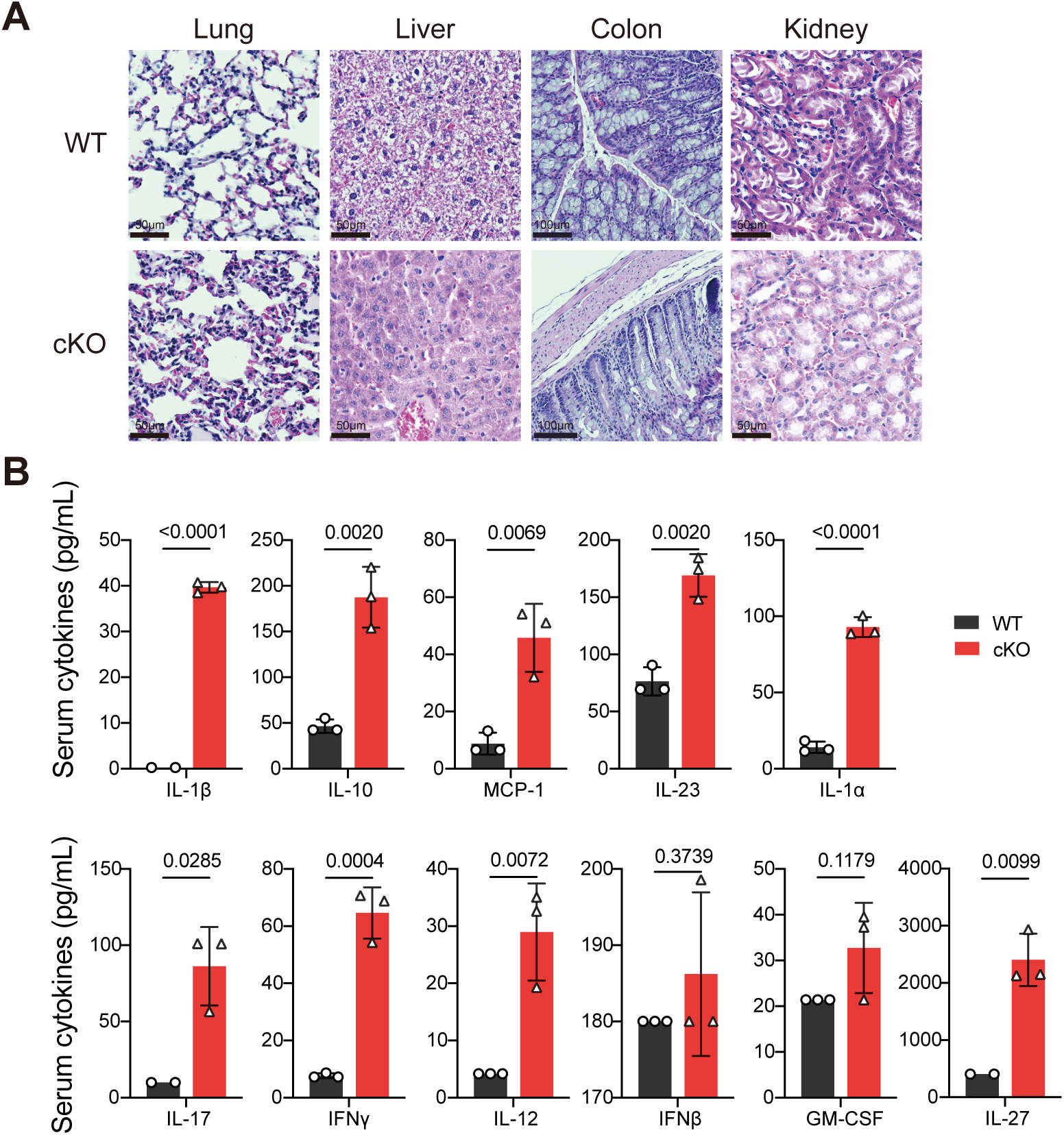
Increased cell infiltration and inflammation in different organs in Foxp3^Cre^Tfam^f/f^ mice. **A.** Histology sections stained with H&E from lung, liver, colon, and kidney. **B.** Serum cytokines identified by Legend plex for IL-1β, IL-10, MCP1, IL-23, IL-1α, IL-17, IFNγ, IL-12, IFNβ, GM-CSF, and IL-27 with estimated values displayed. Data represented as mean ± SD. Statistical test used: t-test, WT (n=5-10; M/F) and cKO mice (n=5-10, M/F).

**Supplementary Fig. 6:**
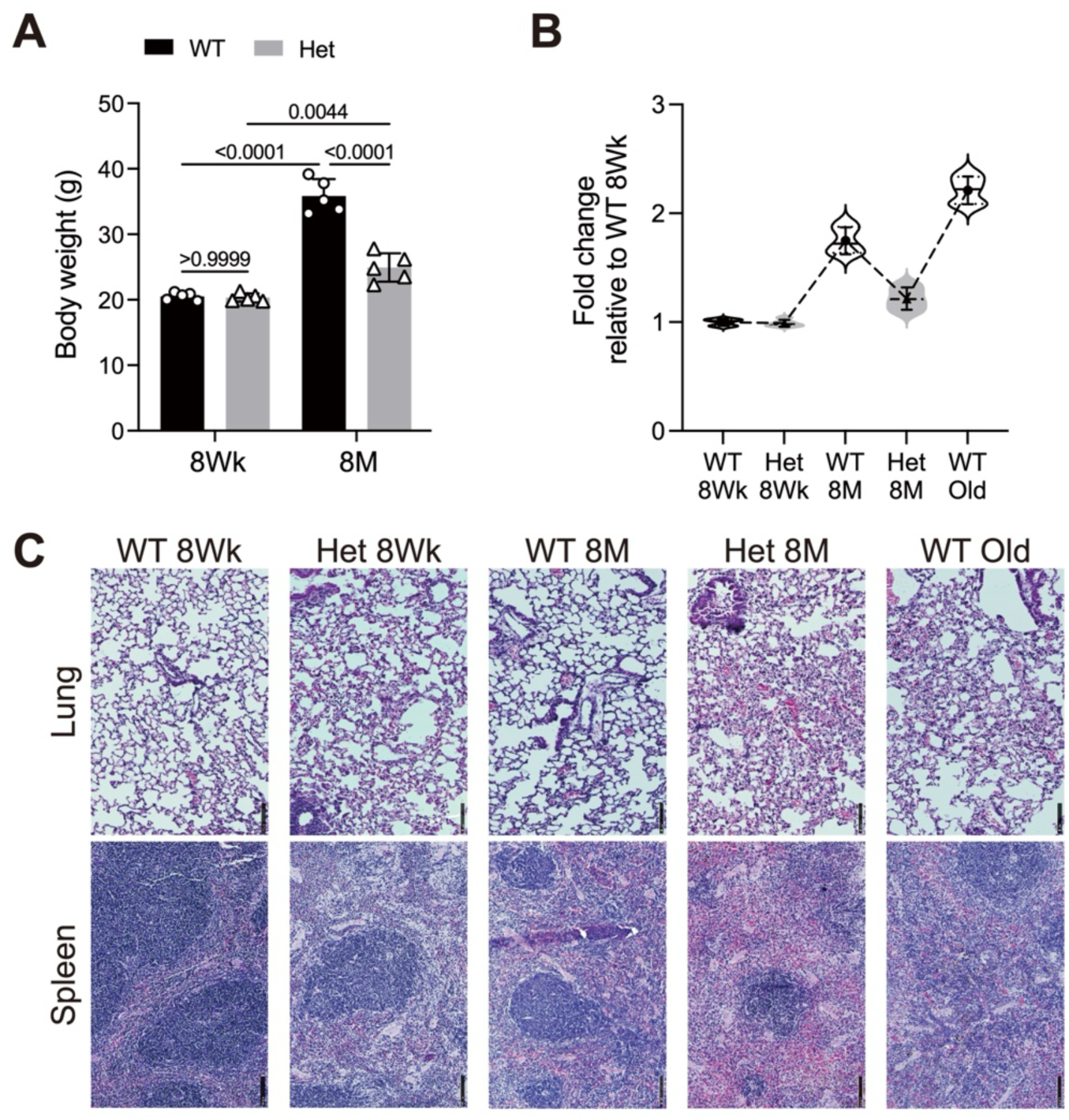
Restoration of mitochondrial function leads to mitigate pathologies. **A-B.** Body weight (**C**) and fold change (**D**) of young and old WT and cKO mice. Data represented as mean ± SD. Statistical test used: two-way ANOVA with post-hoc test (C). WT (n=5; M/F) and Het mice (n=5, M/F).

**Supplementary Table 1:** Gene Set Enrichment Analysis (GSEA) of KEGG pathway between cKO and WT for all cell types.

**Supplementary Table 2:** Gene Set Enrichment Analysis (GSEA) of KEGG pathway between cKO and WT for CD4 subsets.

**Supplementary Table 3:** Functional enrichment analysis of KEGG pathway for four clusters.

**Supplementary Table 4:** Functional enrichment analysis of KEGG pathway for Monocle states.

**Supplementary Table 5:** Gene Set Enrichment Analysis (GSEA) of KEGG pathway between cKO and WT from Brain, Spleen and Skin RNA-seq.

**Supplementary Table 6:** Primer sequences for RT-qPCR, antibodies for Western blots and flow cytometry, and other reagents.

## Acknowledgments

We thank Kristian Herman, Reet Goyal, Risha Mathur, Leia Laurer, Sofie Robinson, and Harpreet Singh for assistance with cell culture, mouse experiments, and general laboratory support. Technical support was provided by Dr. Bony De Kumar and Damien Parrello (Genomics Core), Steve Anderson and Subha Nookala (Flow Cytometry Core), Neeta Adhikari and Donna Laturnus (Histology Core), Sarah Abrahamson (Imaging Core), Ellen Olson and Dr. Collin Combs (Behavior Core), and Dr. Brett McGregor (Computational Data Analysis Core) at the University of North Dakota.

## Contributions

R.M. conceived and designed the study. K.G., R.M., J.G., and Z.W. analyzed and interpreted the scRNA-seq and RNA-seq data. T.D., Z.W., J.G., B.B., H.M., A.L., R.P., M.R., and S.J. performed the experiments. R.M., K.G., J.G., and Z.W. prepared the figures and wrote the manuscript. R.M., J. G., H.C., K.G., N.K., J.H., Z.W., D.J., H.B., M.G., and A.T. critically reviewed and revised the manuscript. All authors contributed to discussions and approved the final version.

